# Abnormal ventricular wall patterning precedes and drives MYBPC3 hypertrophic cardiomyopathy

**DOI:** 10.64898/2026.03.25.714341

**Authors:** Alejandro Salguero-Jiménez, Alba Pau-Navalón, Marcos Siguero-Álvarez, Carlos Relaño-Rupérez, Javier Santos-Cantador, María Sabater-Molina, Xiaoxi Luo, Laura Lalaguna, Laura Sen-Martín, Daniel Martín Pérez, Abel Galicia Martín, Bin Zhou, Juan Antonio Bernal Rodríguez, Fátima Sánchez-Cabo, Enrique Lara-Pezzi, Jorge Alegre-Cebollada, Juan R. Gimeno-Blanes, Donal MacGrogan, José Luis de la Pompa

## Abstract

**BACKGROUND:** Excessive trabeculations and myocardial crypts are recurrent features across cardiomyopathies, yet their developmental origins and clinical significance remain poorly defined. To reveal the link between cardiac morphogenesis and disease, we generated humanized mouse models carrying patient-derived *MYBPC3* frameshift mutations associated with overlapping hypertrophic cardiomyopathy (HCM) and left ventricular non-compaction (LVNC).

**METHODS:** We applied CRISPR-Cas9 to introduce distinct *MYBPC3* frameshift alleles into the mouse genome and performed comprehensive phenotypic and transcriptomic profiling from fetal life through adulthood.

**RESULTS:** Adult homozygous *Mybpc3* frameshift mutant mice like humans displayed hallmark HCM; however, without LVNC. Fetal and neonatal mutant hearts exhibited markedly enlarged ventricular trabeculae and crypts that progressed postnatally into the observed adult hypertrophy. Transcriptomic analysis revealed stage-specific dysregulation of oxidative metabolism, nonsense-mediated decay (NMD), and cell cycle pathways, peaking at postnatal days 1 and 7, indicating that these stages represent critical time points in disease onset. The persistent NMD signature, also observed in phenotype-negative heterozygotes, suggests a compensatory stress response. Enlarged trabeculae exhibited 2-fold increased trabecular cardiomyocyte proliferation, reversing the normal compact–trabecular proliferative gradient and leading to impaired ventricular compaction in neonates. Hey2^CreERT2^ lineage tracing demonstrated invasion of Hey2^+^ compact cardiomyocytes into the trabeculae and ectopic trabecular expression of the Prdm16 transcription factor, indicating defective ventricular wall patterning and maturation. Postnatally, Hey2^+^-derived cardiomyocytes became restricted to the outer/compact myocardium in mutants, while the inner/trabecular myocardium underwent accelerated hypertrophy concurrent with *Prdm16* downregulation. Mice with a *Mybpc3* missense variant also exhibited Hey2^+^ myocardial lineage expansion into trabeculae but no increased proliferation, implicating additional mechanisms beyond Hey2 regulation. Postnatal Prdm16 restoration, via transgenic expression in Mybpc3-null mice effectively attenuated hypertrophy, establishing a causal link between Mybpc3 loss, Prdm16 decline, and pathological remodeling.

**CONCLUSIONS:** Mybpc3 governs ventricular wall maturation by regulating cardiomyocyte proliferation, patterning, and maturation, partly via Prdm16. Disruption of these developmental programs precedes and drives adult HCM, highlighting a developmental role for sarcomeric proteins, and revealing postnatal Prdm16 modulation as an antihypertrophic therapeutic strategy.

## INTRODUCTION

Genetic cardiomyopathies are inherited disorders of the heart muscle caused by mutations that disrupt myocardial structure or function, often resulting in heart failure, arrhythmias, or sudden cardiac death. Mutations in sarcomere genes, especially those encoding contractile structures, are a leading cause across populations. Such mutations are associated with a broad spectrum of clinical phenotypes, including hypertrophic (HCM), dilated (DCM), restrictive (RCM), and left ventricular non-compaction (LVNC) cardiomyopathies ^1^. Mounting evidence reveals significant genetic and phenotypic overlap among these subtypes, complicating accurate diagnosis and clinical management. ^2–4^ The coexistence of HCM and LVNC within individual patients underscores this complexity. HCM is defined by asymmetric left ventricular hypertrophy, myocyte disarray, and interstitial fibrosis, typically in the absence of increased afterload. ^5^ In contrast, LVNC is defined by excessive trabeculation combined with a bilayered myocardium, featuring a compacted epicardial layer alongside a prominent non-compacted endocardial layer, reflecting incomplete myocardial compaction during embryogenesis. ^4, 6^ Clinically, LVNC displays a highly variable severity, ranging from asymptomatic presentation to severe heart failure or sudden death. Patients that present with a mixed HCM-LVNC phenotype, in which hypertrophy and trabeculation coexist, has been associated with a more severe and unpredictable clinical trajectory ^7–12^. This variability suggests that disease expression is shaped by a combination of genetic, epigenetic, developmental, and environmental modifiers.

*MYBPC3*, which encodes cardiac myosin-binding protein C, is essential for sarcomere structural integrity and normal contractile function. ^13–15^ Pathogenic variants in *MYBPC3*, including truncating and missense mutations, represent the most common genetic cause of HCM and have also been implicated in other inherited cardiomyopathies such as DCM, RCM, and LVNC. ^3, 9, 16, 17^ These *MYBPC3* mutations typically lead to haploinsufficiency due to protein truncation or disrupt sarcomere function through missense variants that affect protein–protein interactions or impair myofilament incorporation. ^18, 19^ *MYBPC3* variants are associated with wide-ranging clinical outcomes, from asymptomatic carriers to individuals with advanced heart failure or sudden cardiac death. ^20^ Notably, even in the absence of overt hypertrophy, mutation carriers may exhibit subtle myocardial abnormalities such as crypts, increased trabeculation, or early systolic dysfunction, suggesting subclinical disease. ^16, 21^ While longitudinal data on the progression of the HCM-LVNC phenotype remain limited, the observed clinical heterogeneity supports the influence of modifier genes and developmental pathways. Variants affecting cardiomyocyte proliferation, trabecular remodeling, and hypertrophic signaling may determine whether early morphological defects remain subclinical or evolve into overt non-compaction with secondary hypertrophy and fibrosis.

In this study, we generated two humanized mouse models harboring patient-derived *Mybpc3* frameshift mutations associated with overlapping HCM–LVNC phenotypes to determine whether perturbations in cardiac development contribute to adult cardiomyopathy. By integrating molecular, developmental, cardiomyocyte lineage-tracing, transcriptomic, and functional rescue analyses from fetal stages through adulthood, we show that both mutations lead to premature transcript termination and loss of protein expression. Mybpc3 deficiency initially disrupts ventricular wall patterning and maturation, resulting in excessive trabeculation and myocardial crypt formation during fetal and early postnatal stages. These early abnormalities are associated with altered cardiomyocyte proliferative gradients, dysregulated metabolic and stress-response pathways, as well as defective specification of compact and trabecular myocardial programs. Although these morphological defects partially resolve with maturation, they are followed by accelerated postnatal hypertrophy and overt HCM, establishing a developmental continuum linking transient non-compaction–like/hypertrabeculation phenotypes to adult cardiomyopathy. Finally, we identify Prdm16 as a key downstream effector of Mybpc3-dependent ventricular maturation and demonstrate that postnatal Prdm16 restoration attenuates the observed hypertrophic remodeling within Mybpc3 nonsense mutants. Our findings establish a developmental continuum linking transient non-compaction–like phenotypes to adult cardiomyopathy and identify postnatal myocardial maturation pathways as potential targets for early diagnosis and therapeutic intervention in genetic heart disease.

## METHODS

### Data availability

The authors declare that all data that support the findings of this study are available within the article and its Supplemental Material. The data, analytical methods, and study materials will be available to other researchers for purposes of reproducing the results or replicating the procedure. The RNA-seq data are deposited in the NCBI GEO database under private accession number **qjcpoieczbghfyv.**

### Ethics and DNA collection

Clinical and genetic studies were conducted in accordance with the Declaration of Helsinki and approved by the Ethics Committee of Clinical Research of Hospital Universitario Virgen de la Arrixaca (218/C/2020). Written informed consent was obtained from all participants. Patients underwent clinical evaluation including ECG and two-dimensional and Doppler echocardiography. Pedigrees were constructed and first-degree relatives were screened using the same protocol. Genomic DNA was extracted from 1 mL of EDTA-treated blood using the DNeasy Blood & Tissue Kit (Qiagen, 69506). Sarcomeric genes *MYBPC3, MYH7, TNNT2, TNNI3, TPM1,* and *GLA* were sequenced in probands to identify candidate disease-causing mutations.

### Mice

Established mouse strains used in this study were R26^mTmG 22^, Hey2^CreERT2 23^, Tnnt2^Cre 24^, Myh6^MerCreMer 25^ and Mybpc3 p.Arg502Trp (R502W). ^26^ The Mybpc3 p.Pro109Serfs*12, Mybpc3 p.Arg887Alafs*160 and Rosa26^Prdm16^ mouse lines were generated as part of this study: (see next section). Animal studies were approved by the CNIC Animal Experimentation Ethics Committee and by the Community of Madrid (Ref. PROEX 054.6/25). All animal procedures conformed to EU Directive 2010/63EU and Recommendation 2007/526/EC regarding the protection of animals used for experimental and other scientific purposes, enacted in Spanish law under Real Decreto 118/2021 (modification on Real Decreto 53/2013) and Law 32/2007.

### Generation of Mybpc3 p.P109Sfs*12 and Mybpc3 p.R887Afs*160 mouse lines

Knock-in mouse lines harboring the mutations found in HCM and LVNC patients were generated using CRISPR-Cas9 technology. Specific sequences for CRISPR RNAs (crRNAs) were designed using Breaking-Cas (http://bioinfogp.cnb.csic.es/tools/breakingcas/), Sanger Institute’s HTGT WGE (https://www.sanger.ac.uk/htgt/wge/) and CRISPOR-TEFOR (http://crispor.tefor.net/crispor.py) online tools. crRNA sequences with less off-targets and higher efficiency scores as close to the nucleotide of the intended mutation as possible were selected. Synthetic crRNA and transactivating CRISPR RNA (tracrRNA) were used (AltR CRISPR-Cas9 system, IDT) in combination with AltR *S. p*. Cas9 Nuclease (1081058, IDT). Single stranded oligodeoxynucleotides (ssODN) were designed to insert the intended mutations and, in the p.Arg887Alafs*160 line, silent variants disrupting the protospacer adjacent motif (PAM) sequence and allowing polymerase chain reaction (PCR) genotyping. Please see the *Major Resources Table* for editing reagents.

### Generation of *Rosa26^Prdm16^* mouse line

A full-length mouse *Prdm16* cDNA cloned into pcDNA3.1 was obtained from addgene (#15503). The plasmid was digested with *SpeI* and *XbaI* to excise *Prdm16* cDNA. The digestion product was cloned into a *pBigT-IRESeGFP* plasmid previously generated by cloning a *SalI IRES-eGFP* fragment into the *Xho*I site of *pBigT*. The resulting *loxP*-*PGK-Neo*-3X-STOP-*loxP*-*Prdm16*-*IRES-eGFP* fragment was cloned into the *Pac*I and *Asc*I sites of the *pROSA26PA* plasmid (Figure S18). Gene targeting of this construct was performed in G4 mouse embryonic stem cells (mESCs) and confirmed by Southern blotting with external 5′ and 3′ hybridization probes (Figure S18). Mice were generated by injecting targeted cells into B6CRL blastocysts to generate chimeras that were then analyzed for germline transmission. The selected animals were backcrossed to the C57BL/6 background. Please see the *Major Resources Table* for genotyping.

*Supplemental Methods* are detailed in the Supplemental Material

## RESULTS

### *MYBPC3* frameshift mutations drive mixed HCM/LVNC phenotypes and define null alleles in mice

We identified the MYBPC3 p.Pro108Serfs*9 ^27^ and MYBPC3 p.Arg891Alafs*160 ^28^ variants in cardiomyopathy families designated as families 1 and 2. Variant carriers exhibited HCM, LVNC, or a combined HCM/LVNC phenotype (Figure 1A,B). Representative cardiac magnetic resonance imaging (CMRI) is shown in Figure 1C-F’, including an unaffected individual from family 1 (III-2, Figure 1C,C’) and mutation carriers with isolated LVNC (III-6, Figure 1D,D’) or mixed HCM/LVNC (III-8, Figure 1E,E’). In family 1, the proband was diagnosed at 42 years during evaluation of cardiac symptoms. Echocardiography revealed left ventricular hypertrophy (LVH), which was confirmed by CMRI. Genetic study identified a pathogenic *MYBPC3* variant (p.Pro108Alafs*9). Eight additional relatives were carriers. Among carriers, 7 out of 8 exhibited HCM and 4/8 showed LVNC, including 3 individuals with overlapping HCM/LVNC features. Hypertrophy was asymmetrical septal in six individuals and concentric in one. Two patients had a significant left ventricular outflow tract gradient (>30 mmHg). Left ventricular ejection fraction was preserved in all carriers. Arrhythmias (atrial fibrillation) occurred in two individuals. Three deaths were recorded: one sudden cardiac death (male, 50 years), one stroke-related death (female, 80 years), and one unrelated death (female, 76 years).

**Figure 1.**
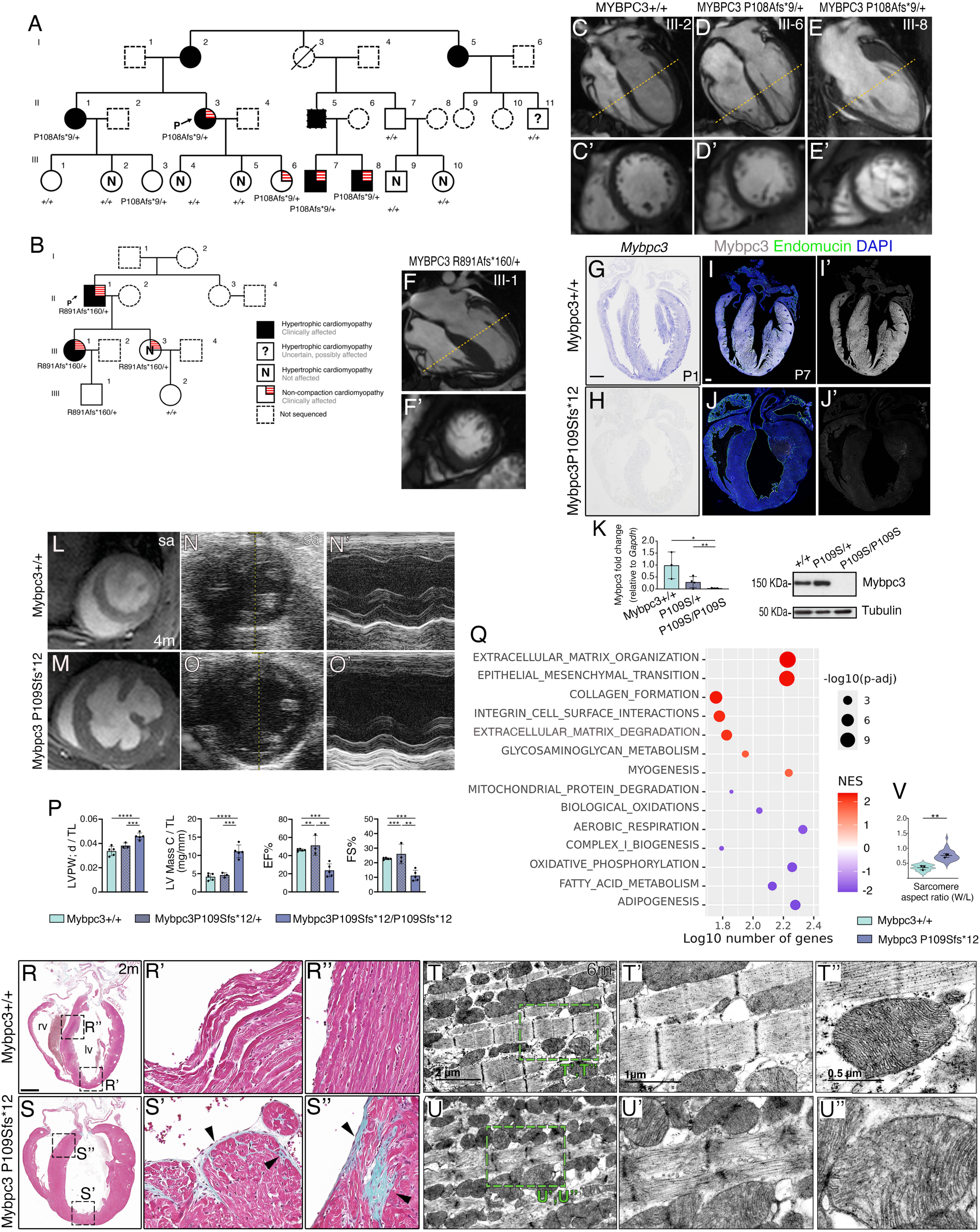
Patient-derived *MYBPC3* mutations cause postnatal hypertrophic remodeling and impaired metabolic maturation in mice. (**A,B**) Pedigrees of two unrelated families carrying *MYBPC3* nonsense variants associated with HCM with variable expressivity and overlapping LVNC. Filled symbols indicate affected individuals; red outlines denote individuals with LVNC or mixed phenotypes; N indicates genotype-negative or unaffected individuals. (**C–E**’) Representative CMRI images from non-affected individuals (**C**) and heterozygous MYBPC3^P108Afs*9^ mutation carriers showing trabeculations (**D,E**). The yellow dashed lines indicate levels of short-axis views (**C’-E’**) highlighting excessive trabeculations (**D’,E’**). (**F,F’**) CMRI of a heterozygous MYBPC3^R891Afs*160/+^ mutation carrier showing prominent trabeculae. (**G,H**) *Mybpc3* ISH in whole-heart sections of control (**G**) and homozygous Mybpc3 p.Pro109Serfs*12 mice at P1 (**H**). (**I,J**) Mybpc3 IF in sections of P7 control (**I**) and mutant hearts (**J**). Mybpc3, grey; Endomucin, green, and DAPI (nuclei), blue. (**I’,J’)** Mybpc3 alone. Scale bar, 250μm. (n=3, ≥4 sections per heart). (**K**) qRT-PCR and WB of P1 control, heterozygous and homozygous mutant Mybpc3 p.Pro109Serfs*12 hearts. (n=3 Mybpc3+/+ and Mybpc3P109Afs*12/+ hearts, and n=4 Mybpc3P109Afs*12/P109Afs*12 hearts). Data are represented as means ± SD. Statistical significance was determined by unpaired Student’s t-test (* p-value <0.05, \*\**p*-value < 0.01). (**L–O’**) Echocardiographic assessment of 4-month-old adult hearts demonstrating concentric hypertrophy in homozygous Mybpc3 p.Pro109Serfs*12 mutant mice, including representative parasternal short-axis views and M-mode recordings (**N’,O’**). (**P**) Quantification of left ventricular wall thickness, mass, ejection fraction (EF), and fractional shortening (FS) across genotypes. (n=5 Mybpc3+/+ and Mybpc3 P109Afs*12/P109Afs*12 animals, n=3 Mybpc3 P109Afs*12/+ animals). Data are represented as means ± SD. Statistical significance was determined by unpaired Student’s t-test (\*\**p*-value < 0.01, \*\*\**p*-value < 0.001, \*\*\*\**p*-value < 0.0001). (**Q**) Gene set enrichment analysis (GSEA) for homozygous Mybpc3 p.Pro109Serfs*12 adult ventricular transcriptome vs control against Hallmark and Reactome gene sets. The bubble plot represents enrichment data for the top gene sets by false discovery rate (FDR). Scale bar indicates normalized enrichment score (NES). Gene count is the number of genes at the intersection between the complete collection of genes used as input for the analysis and the complete list of genes included in the gene set, according to the corresponding database. Longitudinal axis shows the decimal logarithm of the gene count. (**R–S’’**) Histological analysis of adult ventricles (Masson’s trichrome and H&E staining) showing myocardial disarray and interstitial fibrosis in 2-month-old Mybpc3 p.Pro109Serfs*12 mutant hearts (arrowheads). Scale bar, 1mm. (**T–U’**) Transmission electron microscopy (TEM) revealing sarcomeric disorganization, mitochondrial abnormalities, and altered myofibrillar alignment in 6-month-old Mybpc3 p.Pro109Serfs*12 mutant cardiomyocytes compared with controls. (**V**) Sarcomere aspect ratio quantification. (n=3, ≥4 sections per heart). Data are represented as means ± SD. Statistical significance was determined by unpaired Student’s t-test (\**p*-value < 0.05; \*\**p*-value < 0.01).

In family 2, the proband was diagnosed incidentally at 44 years of age. Echocardiography revealed left ventricular hypertrophy (LVH), and Holter monitoring detected non-sustained ventricular tachycardia. Genetic analysis identified a pathogenic *MYBPC3* variant (p.Arg891Alafs*160). An implantable cardioverter-defibrillator was recommended for primary prevention of sudden cardiac death. Three additional relatives were identified as variant carriers. Among the four carriers, 2/4 exhibited HCM and 2/4 showed LVNC, including one individual with overlapping HCM/LVNC features (III-1, Figure 1F,F’). Hypertrophy was asymmetric septal in one case and concentric in one. None of the carriers had a significant left ventricular outflow tract gradient (>30 mmHg). Left ventricular ejection fraction was preserved in all individuals. No arrhythmias or major adverse clinical events were observed during follow-up. Clinical characteristics and outcomes of carriers from both families are summarized in Table S1.

To investigate the molecular mechanisms underlying this striking phenotypic diversity, we generated mouse models carrying orthologous *Mybpc3* variants using CRISPR-Cas9 gene editing. The Mybpc3 p.Pro109Serfs*12 mutation was generated by a single-nucleotide insertion in exon 3 that substituted proline 109 with serine and caused a frameshift that introduced a premature termination codon (PTC) 12 amino acids downstream (Figure S1A-C). This frameshift was predicted to yield a 40–amino acid truncated peptide retaining only the cardiac-specific N-terminal Ig-like C0 domain ^29^. In situ hybridization (ISH) and qRT-PCR analyses confirmed robust *Mybpc3* mRNA expression in control postnatal day 1 (P1) hearts, whereas transcript was undetectable in homozygous Mybpc3 p.Pro109Serfs*12 hearts (Figure 1G,H,K). Consistently, immunofluorescence (IF) and Western blot (WB) analyses, using a monoclonal antibody recognizing the first 120 amino acids of Mybpc3 ^30^, revealed no detectable Mybpc3 protein in homozygous Mybpc3 p.Pro109Serfs*12 P7 hearts (Figure 1I-J’,K). Interestingly, heterozygous Mybpc3 p.Pro109Serfs*12 mice showed normal Mybpc3 protein levels despite a marked reduction in *Mybpc3* mRNA expression (Figure 1K), suggesting the presence of a post-transcriptional compensatory mechanism in the heterozygous state.

The Mybpc3 p.Arg887Alafs*160 allele was generated by a single-nucleotide insertion in exon 23, resulting in a frameshift within a conserved region (human Arg891; mouse Arg887). This introduces a PTC 160 amino acids downstream, producing a 1051–amino acid truncated protein lacking the C-terminal domain (Figure S1A,D,E). ISH at postnatal day 1 (P1) showed *Mybpc3* mRNA expression in the cardiac ventricles of control mice, which was markedly reduced in homozygous Mybpc3 p.Arg887Alafs*160 hearts (Figure S2A,B). IF analyses using the anti-Mybpc3 antibody described above, confirmed robust Mybpc3 protein expression in control hearts, but absence of detectable protein in homozygous mutants (Figure S2C-D’). qRT-PCR analysis revealed significantly reduced expression in both heterozygous and homozygous Mybpc3 p.Arg887Alafs*160 mutants compared to *Mybpc3* transcript levels in control mice (Figure S2E). Despite the reduction in mRNA, WB showed normal levels of MYBPC3 protein in heterozygous hearts, while homozygous hearts lacked detectable protein (Figure S2F).

Both the amino- Mybpc3 p.Pro109Serfs*12 and more carboxy-terminal Mybpc3 p.Arg887Alafs*160 mutations, although located at opposite ends of Mybpc3, abolish detectable protein expression in homozygous mutants, confirming both frameshift mutations as null alleles. In contrast, heterozygotes mice retain normal Mybpc3 protein levels (Figure 1K, Figure S2F) despite reduced mRNA, suggesting post-transcriptional compensation. These findings have also been observed in human patient samples and hiPSC harboring *MYBPC3* inactivating mutations.^19, 31–34^

### Cardiac dysfunction, fibrosis, and sarcomere disarray in Mybpc3-null hearts

We evaluated cardiac structure and function in both homozygous Mybpc3 p.Pro109Serfs*12 and Mybpc3 p.Arg887Alafs*160 4 months-old adult mice by CMRI and echocardiography (Figure 1L-O’; Figure S4A-I). We observed an overall enlargement of the heart in both homozygous mutant mice, with increased left ventricular posterior wall (LVPW) thickness, LV volume and mass (Figure 1L-O’,P; Figure S4J-M), typical of HCM. Cardiac function was also impaired in these mutant mice, with reduced ejection fraction (EF) and fractional shortening (FS) (Figure 1P; Figure S4N,O). These mutations recapitulate features observed in human non-obstructive HCM echocardiography (increased LV wall thickness and reduced EF), and mirror morphological and functional deficits linked to Mybpc3 deficiency in mouse models. ^13, 35^

We next employed bulk RNA-sequencing (RNA-seq) to investigate the transcriptional changes in adult Mybpc3 p.Pro109Serfs*12 hearts (Figure 1Q; Tables S2 and S3). Gene Set Enrichment Analysis (GSEA) revealed prominent enrichment of pathways associated with extracellular matrix (ECM) organization, including collagen biosynthesis, degradation, and glycosaminoglycan metabolism, indicating active ECM remodeling and dysregulated collagen turnover (Figure 1Q, Figure S5, Table S3). Consistent with this molecular signature, Masson’s Trichrome staining in 2-month-old Mybpc3 p.Pro109Serfs*12 adult hearts showed focal subendocardial fibrosis in the left ventricle (Figure 1R,R’,S,S’), and interstitial fibrosis in the ventricles and septum (Figure 1R’’,S’’). Similar fibrotic patterns were observed in Mybpc3 p.Arg887Alafs*160 mice (Figure S4P-Q’’) mirroring a hallmark feature of HCM: excessive collagen deposition. ^36^ Mutant hearts also displayed a robust myogenesis-related transcriptional signature (Figure 1Q), consistent with pathological cardiac hypertrophy. ^37^ In addition, an NMD-associated gene signature was enriched, aligning with chronic activation of RNA surveillance pathways in HCM driven by the *Mybpc3* mutation (Figure S5). ^34, 38, 39^ In contrast, gene signatures associated with oxidative phosphorylation, fatty acid metabolism, and mitochondrial protein degradation, were depleted (Figure 1Q, Figure S5), indicating impaired mitochondrial function and metabolic remodeling. Adipogenesis, which contributes to energy balance and offers cardioprotective effects in the human heart, ^40^ was also negatively enriched, suggesting a loss of adaptive metabolic buffering capacity.

Transmission electron microscopy (TEM) analysis of 6-month-old adult homozygous Mybpc3 p.Pro109Serfs*12 hearts revealed marked ultrastructural abnormalities, including disrupted and misaligned sarcomeres, irregular Z-lines, and increased interstitial space in homozygous mutant animals (Figure 1T-U’,V). These changes have been implicated in reducing actin–myosin overlap and active force generation. ^41^ Furthermore, the mitochondria appear enlarged and variably shaped, with regions of disorganized cristae, consistent with mitochondrial remodeling in the Mybpc3-deficient myocardium (Figure 1T’’,U’’). These findings reveal structural abnormalities at both the sarcomere and mitochondrial levels, mirroring the cellular and molecular alterations characteristic of HCM.

In summary, adult Mybpc3 p.Pro109Serfs*12 and Mybpc3 p.Arg887Alafs*160 mutants faithfully recapitulate key pathological features of HCM, including cardiomyocyte hypertrophy, myocardial fibrosis, sarcomere disarray, and impaired energy metabolism. However, they do not exhibit overt structural abnormalities such as the LVNC/hypertrabeculation phenotype observed in human carriers.

### Hypertrabeculated, fetal-like hearts progress to pathological hypertrophy in *Mybpc3* mutants

To get a deeper insight into the cardiac phenotype of these *Mybpc3* mutants we analyzed heart morphology at postnatal day 1 (P1), when ventricular development is well advanced (Figure 2A-A’’). Homozygous Mybpc3 p.Pro109Serfs*12 (Figure 2B-B’’) and Mybpc3 p.Arg887Alafs*160 (Figure 2C-C’’) mutant hearts displayed profound structural abnormalities, predominantly affecting the left ventricle. This included chamber dilation, markedly reduced compact myocardium (CM) thickness, and elongated trabecular myocardium (TM) with large apical and free-wall trabeculations extending into the ventricular lumen (Figure 2B’-C’’,G). The large trabeculae of homozyogous P1 Mybpc3 p.Pro109Serfs*12 and Mybpc3 p.Arg887Alafs*160 mutants, resembled myocardial “crypts” ^42^ and potentially will lead to functional impairments preceding the development of overt HCM. ^35^

**Figure 2.**
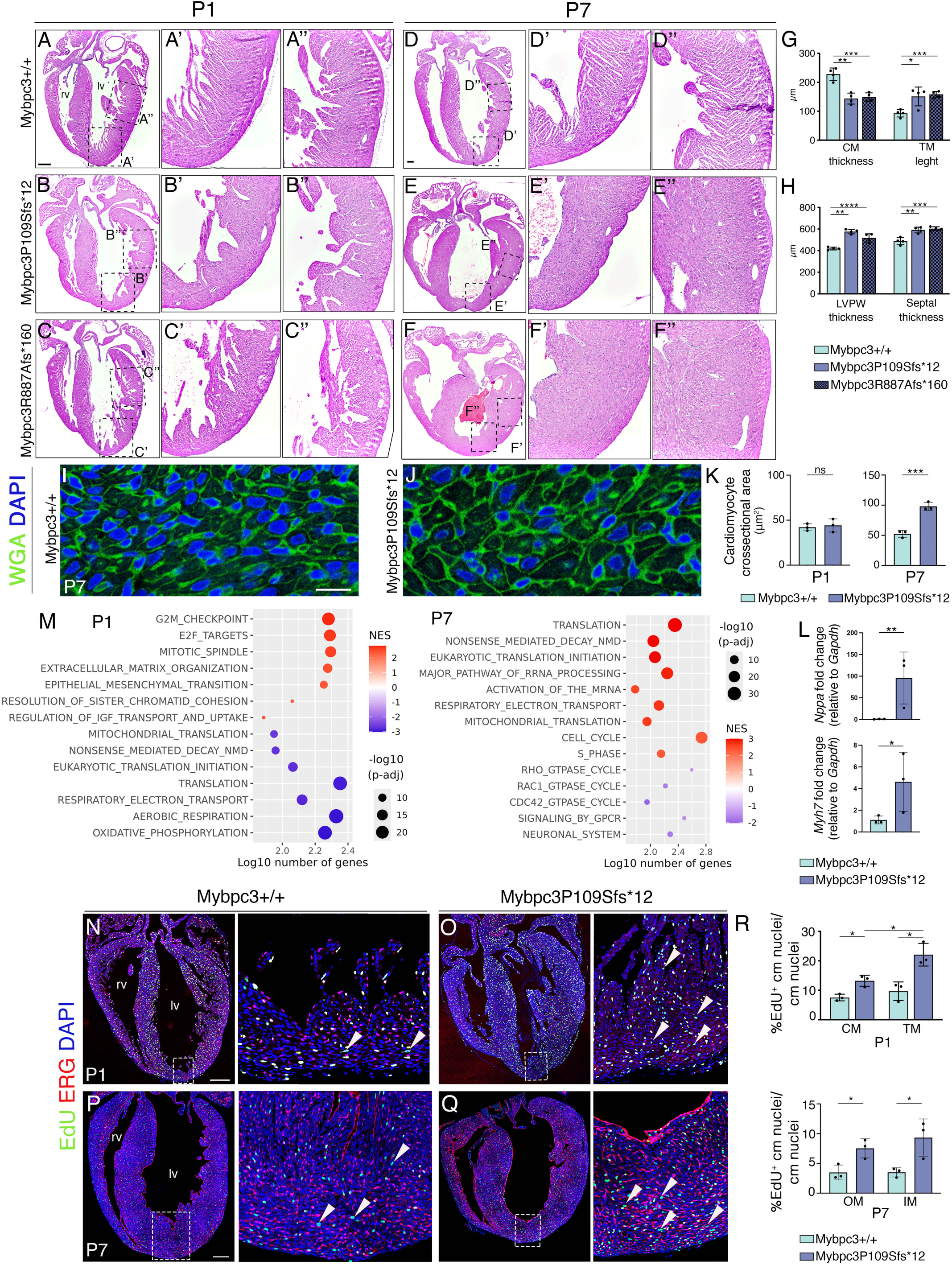
Loss of mouse Mybpc3 drives a neonatal switch from hypertrabeculation/non-compaction to hypertrophy during early postnatal heart maturation. H&E-stained heart sections of control Mybpc3^+/+^, homozygous Mybpc3 p.Pro109Serfs*12 and homozygous Mybpc3 p.Arg887Alafs*160 P1 (**A-C’’**), and P7 (**D-F’’**) neonate mice. Scale bars, 250 μm. lv, left ventricle; rv, right ventricle. (**G**) Analysis of compact myocardium (CM) thickness and trabecular myocardium (TM) length. (n=4, ≥5 sections per heart). Statistical significance was determined by unpaired Student’s t-test (*p-*value* < 0.05; \*\**p*-value < 0.01; \*\*\**p*-value < 0.001). (**H**) Analysis of left ventricle posterior wall (LVPW) and septum thickness. (n=4, ≥5 sections per heart). Statistical significance was determined by unpaired Student’s t-test (\*\**p*-value < 0.01; \*\*\**p*-value < 0.001; ; \*\*\*\**p*-value < 0.0001). (**I,J**) WGA (green) and DAPI (blue) fluorescence staining of P7 control Mybpc3 +/+ and homozygous Mybpc3 p.Pro109Serfs*12 heart sections. Scale bar, 10 μm. (**K**) Cardiomyocyte cell area quantification using WGA staining of P7 Mybpc3^+/+^ and homozygous Mybpc3 p.Pro109Serfs*12 heart sections. (n=4, ≥5 sections per heart). Data are represented as means ± SD. Statistical significance was determined by unpaired Student’s t-test (ns, not significant; \*\*\**p*-value < 0.001). (**L**) qRT-PCR analysis of *Nppa and Myh7* expression (relative to *Gapdh*) in P7 control and homozygous Mybpc3 p.Pro109Serfs*12 hearts (n= 3 hearts). Statistical significance was determined by unpaired Student’s t-test (\**p*-value < 0.05; \*\**p*-value < 0.01). **(M)** GSEA of ventricular tissue transcriptomics from homozygous Mybpc3 p.Pro109Serfs*12 mutant mice compared to control littermates at P1 and P7. Dot plots display top significantly enriched Hallmark and Reactome pathways. Dot size corresponds to the negative logarithm of the False Discovery Rate (FDR) and color to the Normalized Enrichment Score (NES). Longitudinal axis shows the logarithm of the number of genes present in both the dataset and the gene set. (**N-Q**) 5-ethynyl-2′-deoxyuridine (EdU, green) incorporation IF in P1 and P7 control and homozygous Mybpc3 p.Pro109Serfs*12 hearts. Endocardial nuclei are stained with ERG (red) and nuclei counterstained with DAPI (blue). Arrowheads, proliferative EdU^+^ cardiomyocytes. Scale bar, 250 µm. (**R**) Quantification of the percentage of EdU incorporation in control and homozygous Mybpc3 p.Pro109Serfs*12 mutant hearts at P1 and P7. CM, compact myocardium, TM, trabecular myocardium. OM, outer and IM, inner myocardium; cm, cardiomyocyte. (n=3, ≥3 sections per heart). Data are represented as means ± SD. Statistical significance was determined by unpaired Student’s t-test (ns, not significant; \**p*-value < 0.05).

At P7, control heart shows trabeculae still visible and a thick compact myocardium (Figure 2D-D’’). ^43–45^ Homozygous Mybpc3 p.Pro109Serfs*12 and Mybpc3 p.Arg887Alafs*160 mutant mice exhibited enlarged and dilated heart, with increased left ventricular posterior wall (LVPW) and ventricular septum thickness (Figure 2E-F’’,H), indicative of ventricular hypertrophy. We next measured cellular hypertrophy after labeling with wheat germ agglutin (WGA) the membrane of cardiomyocytes in sections of both P1 and P7 hearts. Examination at P1 did not reveal cardiomyocyte enlargement, as WGA staining revealed similar cross-sectional area of Mybpc3 p.Pro109Serfs*12 cardiomyocytes compared to controls (Figure S6A-C). At P7 we did observe increased cardiomyocyte area in Mybpc3 p.Pro109Serfs*12 homozygous mutant mice compared to controls (Figure 2I,J,K; Figure S6D-F). Additionally, P7 mutant hearts exhibited discrete regions within the left ventricle where sarcomeres appeared disorganized (Figure S7). This focal sarcomeric disorganization at P7 is consistent with an early defect in postnatal sarcomere maturation associated with the *Mybpc3* mutation ^13, 46–48^ and may precede the development of overt hypertrophy. The cardiac hypertrophy markers *Natriuretic peptide type A* (*Nppa*) and *beta Myosin heavy chain* (*Myh7*) were also upregulated by qRT-PCR in P7 Mybpc3 p.Pro109Serfs*12 homozygous mutant hearts (Figure 2L). We observed comparable evidence of cardiomyocyte hypertrophy in Mybpc3 p.Arg887Alafs*160 homozygous mutant hearts (Figure S8). Thus, mice harboring the Mybpc3 p.Pro109Serfs*12 and Mybpc3 p.Arg887Alafs*160 mutations identified from cardiomyopathy patients, display at P1 a hypertrabeculation phenotype that by P7 shows both morphological and molecular features of pathological cardiac hypertrophy.

### Disrupted Postnatal Cardiac Maturation in *Mybpc3* Mutants

To better understand the underlying mechanisms, we examined the molecular and cellular processes affected by *Mybpc3* ablation across postnatal cardiac maturation (Figure 2M, Tables S2 and S3, and Figures S9 and S10). RNA-seq analysis of P1 Mybpc3 p.Pro109Serfs*12 mutant ventricles (Figure 2M) revealed profound molecular perturbations associated with ventricular wall dysmorphology (Figure 2A-F’’). Gene set enrichment analysis (GSEA) identified significant enrichment of pathways associated with the cell cycle, including E2F targets and G2/M checkpoint signaling, and the ECM (ECM organization) (Figure 2M, Figure S9). Tgfß signaling, previously linked to LVNC ^49, 50^ was also increased at this stage (Figure S9). In contrast, pathways related to oxidative metabolism (OxPhos, mitochondrial biogenesis, mitophagy) and RNA transcription/translation (including NMD and ubiquitination) were depleted (Figure 2M; Figure S9). These findings indicate a developmentally immature transcriptional profile, consistent with impaired ventricular wall maturation and compromised energy production and proteostasis.

At P7, Mybpc3 p.Pro109Serfs*12 mutants showed significant enrichment of pathways associated with translation (mTORC1 signaling) and rRNA processing (NMD, major pathway of RRNA processing, mRNA splicing, etc.), Myc-regulated targets (V1 and V2), oxidative metabolism (respiratory electron transport chain, mitochondrial translation OxPhos, Glycolysis), and cell cycle activity (E2F targets, G2-M transition, transcriptional regulation by P53,etc.) (Figure 2M; Figure S10). Conversely, only a limited set of pathways were depleted, including signatures linked to cellular behavior (GTPase cycle), GPCR signaling, the ECM organization and cardiac conduction (Figure 2M; Figure S10). This pattern reflects a transcriptional shift toward a hypertrophic molecular signature that is consistent with the emerging cardiac phenotype. In agreement, P1 and P7 Mybpc3 p.Pro109Serfs*12 homozygous hearts showed markedly increased numbers of S-phase cells in CM and especially in TM (>2-fold increase) compared with control (Figure 2N–Q,R; Figure S11), indicating delayed postnatal cardiomyocyte maturation.

Interestingly, Mybpc3 haploinsufficiency alone did not produce any obvious structural or functional abnormalities: both Mybpc3 p.Pro109Serfs*12/+ and Mybpc3 p.Arg887Alafs*160/+ hearts appeared morphologically normal at P1 and P7 (Figure S3A-F’). However, RNA-seq analysis of Mybpc3 p.Pro109Serfs*12/+ heterozygotes revealed significant enrichment of transcription- and translation-related pathways at both P1 and P7, including NMD, with mTORC1 and cell cycle signatures elevated at P1 but not a P7. The ECM-associated pathway was consistently depleted at both stages (Figure S3G). These findings align with previous studies demonstrating that haploinsufficiency can activate maladaptive molecular programs even when residual *Mybpc3* expression is preserved. ^34, 38, 51^

Collectively, our RNA-seq analyses indicate that Mybpc3 deficiency preserves a fetal-like gene expression profile at P1 followed by the emergence of an enhanced hypertrophic transcriptional program at P7.

### Abnormal Hey2⁺ trabecular lineage contribution and ventricular maturation upon Mybpc3 loss

The Mybpc3 p.Pro109Serfs*12 mutants exhibited no overt morphological defects at E15.5: ventricular wall architecture and trabecular-compact myocardium patterning appeared normal (not shown). In contrast, molecular profiling revealed clear significant changes in gene expression during this fetal ventricular wall maturation and compaction stage, well before the onset of observable HCM phenotype (Figure 3A). GSEA identified enrichment of metabolic and growth-related pathways, including cholesterol homeostasis, mTORC1 signaling, MYC targets (V1 and V2), and IGF transport and uptake (Figure 3A; Figure S12), indicating early alterations in cardiomyocyte growth and metabolic regulation. In parallel, pathways related to transcriptional and translational regulation, such as eukaryotic translation initiation, mRNA processing, NMD, and activation of stress pathways (unfolded protein response), were also upregulated (Figure 3A; Figure S12). In addition, we observed pathways associated with tissue structural maturation were negatively enriched in mutant hearts. These included ECM organization, collagen formation, and glycosaminoglycan metabolism, suggesting impaired ECM remodeling during myocardial maturation (Figure 3A; Figure S12). Signaling pathways involved in cardiac morphogenesis and patterning, such as WNT signaling, were likewise downregulated (Figure 3A). Together, these data show that loss of Mybpc3 at E15.5 triggers early transcriptional dysregulation, affecting cardiomyocyte protein homeostasis, growth-related signaling, and metabolic pathways, while concurrently suppressing ECM and structure-related maturation programs. These molecular abnormalities precede overt postnatal phenotypes and indicate that developmental dysregulation begins during fetal ventricular wall maturation upon Mybpc3 loss.

**Figure 3.**
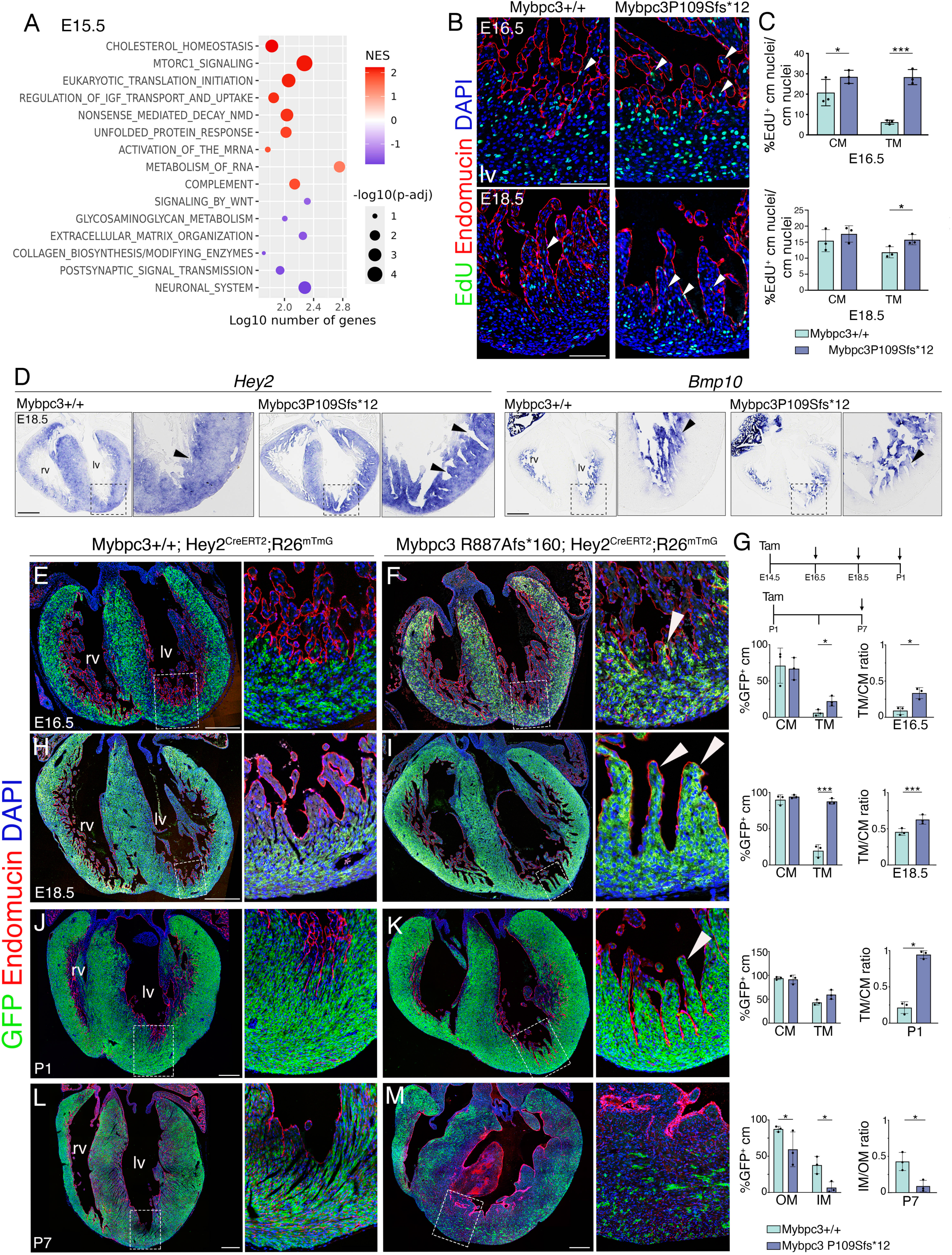
Increased proliferation and contribution of Hey2^+^ cardiomyocytes to the fetal and neonatal trabeculae of Mybpc3-deficient hearts. (**A**) GSEA of differentially expressed genes in ventricular tissue from homozygous E15.5 Mybpc3 p.Pro109Serfs*12 mutant mice compared with control littermates. Dot plots display top significantly enriched Hallmark and Reactome pathways. Dot size corresponds to the negative logarithm of the False Discovery Rate (FDR) and color to the Normalized Enrichment Score (NES). Longitudinal axis shows the logarithm of the number of genes present in both the dataset and the gene set. (**B**) EdU incorporation in E16.5 and E18.5 control and Mybpc3 p.Pro109Serfs*12 homozygous mutant hearts. Endomucin (red), delineates the endocardium, nuclei are counterstained with DAPI (blue). Arrowheads, proliferating cardiomyocytes. Scale bars, 100 μm. lv, left ventricle; rv, right ventricle. (**C**) Quantification of EdU incorporation percentage in E16.5 and E16.5 control and homozygous Mybpc3 p.Pro109Serfs*12 mutant hearts (n=3, ≥3 sections per heart). Data are represented as means ± SD. Statistical significance was determined by unpaired Student’s t-test (ns, not significant; \**p-*value < 0.05; \*\*\**p-*value < 0.001). (**D**) *Hey2* and *Bmp10* ISH in E18.5 control and Mybpc3^P109Sfs*12^ mutant hearts. Scale bars 250 μm. (**E,F, H-M**) GFP (green) IF in E16.5 (**E,F**), E18.5 (**H,I**), P1 (**J,K**) and P7 (**L,M**) Mybpc3^+/+^;Hey2^CreERT2/+^;Rosa26^mTmG/+^ and Mybpc3 p.Arg887Alafs*160;Hey2^CreERT2/+^;Rosa26^mTmG/+^ hearts induced with Tamoxifen (Tam) at E14.5 and analyzed at E16.5, E18.5 or P1, or induced at P1 and analyzed at P7 (**G**). (**F,I,K**,**M**, insets) The endocardium is stained with Endomucin (red) and nuclei with DAPI (blue). Arrowheads indicate Hey2-derived cardiomyocytes (green) in the trabeculae. Scale bars 250 μm. (**G**) Left column, analysis of the Hey2 ^CreERT2^ recombination rate measured by the fraction of GFP^+^ cardiomyocytes (cm) compared to total number of cardiomyocytes in control and homozygous Mybpc3 p.Arg887Alafs*160 mutants. Right column, ratio of TM and CM GFP^+^ cardiomyocytes percentages. (n=3, ≥3 sections per heart). Data are represented as means ± SD. Statistical significance was determined by unpaired Student’s t-test (\**p*-value < 0.05; \*\*\**p*-value < 0.001).

In line with this finding, and consistent with observations at P1 and P7 (Figure 2M-R; Figures S9 & S10), we detected increased trabecular myocardium proliferation in E16.5 and E18.5 Mybpc3 p.Pro109Serfs*12 mutants (Figure 3B,C). The immature CM is more proliferative than the differentiated TM. ^52^ Therefore, we investigated whether the increased TM proliferation observed in Mybpc3-deficient hearts reflects an underlying defect in ventricular patterning. ISH analysis at E18.5 revealed altered expression of the compact marker *Hey2* and trabecular marker *Bmp10*. In Mybpc3 p.Pro109Serfs*12 ventricles, *Hey2*, which is normally restricted to the CM, was abnormally expanded into the TM. Conversely, *Bmp10*, a marker of TM, was reduced, particularly at the base of the trabeculae (Figure 3D). Together, these findings indicate a progressive disruption of ventricular myocardial patterning during fetal heart development.

To investigate the cellular basis of this phenotype, we focused on Hey2^+^ cardiomyocytes, which drive ventricular compaction by expanding from the epicardial toward the endocardial regions of the wall. ^23^ During compaction, TM proliferation declines, and it is gradually replaced by cardiomyocytes derived from the proliferative CM, forming a transient hybrid zone of CM- and TM-derived cells. ^23, 53^ Disruption of Hey2^+^ cardiomyocyte expansion impairs this process leading to LVNC/hypertrabeculation features, underscoring their essential role in ventricular maturation. ^23^ To assess whether this process was altered in homozygous Mybpc3 p.Pro109Serfs*12 and Mybpc3 p.Arg887Alafs*160 mutants, we crossed these mice with the tamoxifen-inducible Hey2^CreERT2^ line ^23^ and the Rosa26^mTmG^ reporter mouse ^22^ and determined what was the contribution of the *Hey2*-lineage to the ventricular wall in these Mybpc3-deficient backgrounds. Initially, Cre recombination was induced at E14.5, and GFP expression was analyzed at E16.5, E18.5, and P1 (Figure 3E-K, Figure S13). To control for variability in tamoxifen delivery, TM recombination was normalized to CM recombination within each embryo (Figure 3G). At E16.5, both *Mybpc3* homozygous mutants showed a significantly higher TM/CM recombination ratio compared to controls (Figure 3E,F,G; Figure S13A,B,I,J). By E18.5, control hearts displayed the expected pattern of Hey2^+^-derived GFP^+^ cardiomyocytes restricted to the CM, with only a few cells observable within the TM (Figure 3H,G), consistent with prior findings in control hearts. ^23^ In contrast, homozygous Mybpc3 p.Pro109Serfs*12 and Mybpc3 p.Arg887Alafs*160 mutants revealed a marked expansion of the Hey2^+^ lineage into the trabecular region, with significantly more GFP^+^ cardiomyocytes compared to controls, especially within Mybpc3 p.Arg887Alafs*160 mutants (Figure 3I,G; Figure S13C,D,I,J). The CM itself contained comparable numbers of GFP^+^ cells in both control and mutant groups (Figure 3G), but the elevated TM/CM recombination ratio confirmed abnormal infiltration of Hey2^+^-lineage derived cells into the trabecular layer. When Cre recombination was analyzed at P1, a similar pattern was observed: mutants retained significantly higher Hey2^+^ contribution to trabeculae (Figure 3J,K,G; Figure S13E,F,I,J). By contrast, tamoxifen induction at P1 resulted in a reduced contribution of Hey2^+^ cells to the inner myocardial wall of P7 mutants (Figure 3L,M,G; Figure S13G-J).

In summary, loss of Mybpc3 function disrupts normal ventricular myocardium maturation. This patterning defect leads to persistent incorporation of Hey2^+^ cardiomyocytes into the TM during fetal and early postnatal development, thereby sustaining excessive trabecular growth, and promoting the progression from hypertrabeculation-to-HCM.

To determine whether our findings in humanized Mybpc3 loss-of-function mice reflect shared developmental mechanisms underlying Mybpc3-related HCM, we analyzed a third humanized model carrying the missense mutation Mybpc3 p.ArgR502Trp mice (R502W), associated with late-onset HCM ^26^. Homozygous Mybpc3 p.ArgR502Trp mutants displayed prominent ventricular crypts at P1 and P7 (Figure S14A-D’) with diminished CM thickness and increased TM/CM ratio (Figure S14E) and increased cardiomyocyte contribution to trabeculae at E18.5 and P1 (Figure S15), without a significant rise in proliferation (Figure S16). These results suggest that distinct *Mybpc3* mutations converge on impaired ventricular patterning and maturation as a common pathogenic mechanism.

### The Mybpc3 p.Pro109Serfs*12 mutation disrupts postnatal transcriptional maturation and metabolic remodeling

To define the biological pathways altered across development, we performed GSEA at E15.5, P1, P7, and adult stages (Figure 4A). Comparative analysis showed that Mybpc3 loss produced the most pronounced transcriptional disturbances at P1 and P7 (Figure 4A), coinciding with the critical postnatal transition from hyperplastic to hypertrophic cardiac growth. ^54–56^ Fetal and Adult stages showed comparatively modest pathway alterations.

**Figure 4.**
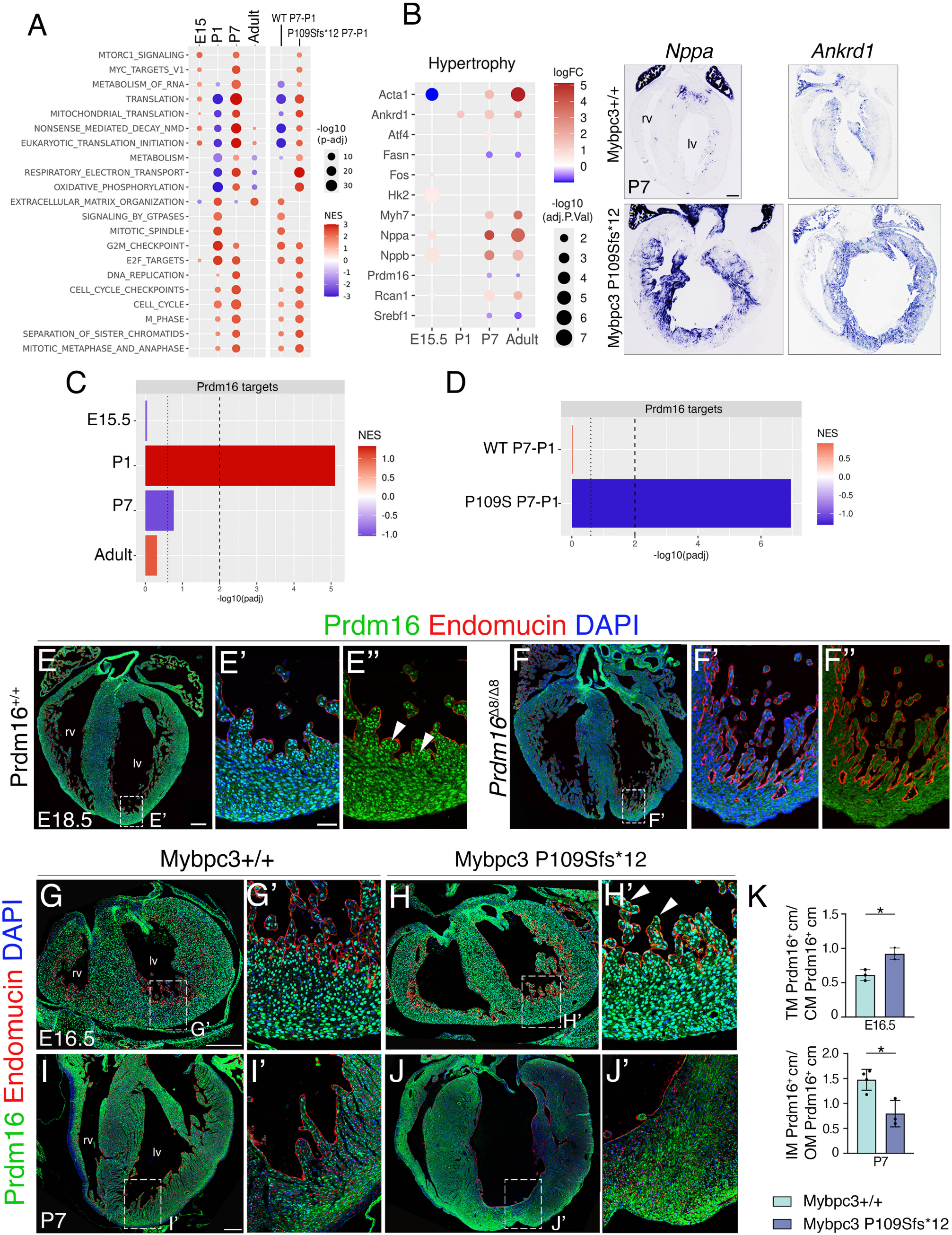
Temporal rewiring of metabolic and cell cycle pathways with dynamic changes in Prdm16 targets and expression in Mybpc3 p.Pro109Serfs*12 mutants. (**A**) Gene set enrichment analysis (GSEA) analysis of ventricular RNA-seq datasets at E15.5, P1, P7 and adulthood comparing homozygous Mybpc3 p.Pro109Serfs*12 and Mybpc3^+/+^ ventricular transcriptome against Hallmark and Reactome gene sets. The first 4 columns show comparisons between mutant and control samples at E15.5, P1, P7 and adult stages. Most significant processes seem to occur at P1 and P7. The mutation causes similar direction changes at P1 and P7 on cell cycle terms (increased in the mutants), but opposite changes in metabolism, NMD and metabolism/proteostasis terms (oxphos, translation) which are decreased at P1 mutants compared to control but increased in the mutants at P7. The last two columns represent the longitudinal comparisons between P1 and P7 in both control and mutant samples. The WT (control) P7-P1 column indicates the developmental change in the transition from P1 to P7 (red, up in P7). NMD and proteostasis (translation etc.) terms are decreased in the controls at P7, but in the mutants, these same processes are increased. Cell cycle terms maintain the same enrichment direction in the mutants but seem to be even more exacerbated at P7. (**B**) Left, bubble plot of differentially expressed genes at P7 in homozygous Mybpc3 p.Pro109Serfs*12 and Mybpc3+/+ hearts highlights upregulation of several genes and downregulation of *Prdm16* within a hypertrophy-associated gene signature. Right, ISH for *Nppa* and *Ankrd1* in control and homozygous Mybpc3 p.Pro109Serfs*12 P7 hearts. 250 μm. lv, left ventricle; rv, right ventricle. ((**C**) Bar plot summarizing the normalized enrichment score (NES) and –log(padj) values for Mybpc3 P109Sfs*12 mutant vs. WT comparisons at E15.5, P1, P7, and Adult. Dotted line: p-value = 0.25. Dashed line: *p*-value = 0.01. **D**) Bar plot summarizing the NES and –log(padj) values for P7 vs. P1 comparisons in control (WT) and Mybpc3 p.Pro109Serfs*12 (P109S) mutant. Dotted line: *p*-value = 0.25. Dashed line: p-value = 0.01. The NES represents the enrichment score after normalization, with higher scores (red) indicating positive enrichment and lower scores (blue) indicating negative enrichment. The adjusted p-value (padj) corresponds to p-values corrected for multiple testing using the Benjamini–Hochberg procedure, which controls the false discovery rate. (**E-F’’**) IF for Prdm16 (green), Endomucin (red), and DAPI (blue) in in ventricular sections of E18.5 wild type (**E-E**’’) and homozygous *Prdm16^Δ8^* mutant hearts. Arrowheads in E’’ point to nuclear Prdm16 expression in trabecular cardiomyocytes. Scale bar 250 μm. Magnified views are shown in (**’**). Scale bar 50 μm. (**G-J’**) IF for Prdm16 in ventricular sections at E16.5 (**G-H’**) and P7 (**I-J’**) reveals expanded Prdm16 signal in E16.5 Mybpc3 p.Pro109Serfs*12 mutant trabeculae (arrowheads, **H’**, inset), and reduced Prdm16 expression at P7 in subendocardial myocardium of mutants (**J’**). Scale bar, 250 μm. (**K**) Quantification of the ratio of trabecular (TM) to compact myocardium (CM) Prdm16 expression in E16.5 hearts (n=3, ≥3 sections per heart), and the ratio of inner myocardium (IM) to outter myocardium (OM) in P7 hearts. (n=4 control and n=3 Mybpc3 p.Pro109Serfs*12, ≥3 sections per heart). Data are represented as means ± SD. Statistical significance was determined by unpaired Student’s t-test (\**p*-value < 0.05).

At P1, mutant hearts showed strong enrichment of cell cycle–associated gene pathways (E2F targets, G2M checkpoint, mitotic spindle, DNA replication, M phase, and cell cycle checkpoints), indicating persistent proliferative activity at birth (Figure 4A). In contrast, pathways tied to oxidative metabolism and proteostasis, oxidative phosphorylation, respiratory electron transport, mitochondrial translation, eukaryotic translation initiation, RNA metabolism, and NMD, were negatively enriched (Figure 4A). Notably, NMD was activated at all stages except P1, consistent with chronic activation of the RNA surveillance program. ECM-related signatures were also altered at this stage, consistent with early postnatal remodeling (Figure 4A). Thus, P1 mutants exhibit a transcriptional profile marked by enhanced proliferation and dampened metabolic and translational programs.

By P7, cell cycle pathways remained positively enriched, with greater magnitude than at P1, suggesting a failure to suppress proliferative programs during normal maturation. Remarkably, metabolic and proteostasis-related pathways reversed directionality relative to P1: oxidative phosphorylation, translation, RNA metabolism, and NMD were now positively enriched in mutants (Figure 4A). This temporal inversion indicates a dysregulated progression of metabolic and RNA-surveillance programs between early (P1) and late (P7) neonatal stages. Cytoskeletal and vesicular transport pathways peaked at P1, whereas ECM-related signatures showed enrichment at P1 and in adults, reflecting postnatal remodeling and early fibrotic progression. In Adults, gene sets associated with ECM organization were enriched, while oxidative metabolism was downregulated (Figure 4A), features consistent with a hypertrophic cardiomyopathy-like state. Together, these results indicate that Mybpc3 deficiency initially delays postnatal maturation (persistent cell cycle activity) and subsequently accelerates hypertrophic remodeling.

To contextualize these changes, we compared P1 and P7 transitions in control and mutant hearts (Figure 4A). In controls, the P1 to P7 transition was marked by strong downregulation of RNA metabolism, translation, mitochondrial translation, and NMD, reflecting the developmental postnatal reduction of biosynthetic programs (Figure 4A, WT (control) P7-P1). In Mybpc3 p.Pro109Serfs*12 hearts (Figure 4A, P109Sfs*12 P7-P1), these same pathways were upregulated, indicating a failure to extinguish early biosynthetic activity. ECM organization and GTPase signaling were positively enriched in WT maturation (WT P7-P1) but showed weaker induction in mutants (P109Sfs*12 P7-P1), indicating impaired structural remodeling (Figure 4A). Cell cycle–related pathways increased between P1 and P7 in controls but were more strongly activated in Mybpc3 p.Pro109Serfs*12 mutants, consistent with sustained proliferative activity. Collectively, these data demonstrate that the Mybpc3 p.Pro109Serfs*12 mutation disrupts the early postnatal transcriptional transition by (i) maintaining cell cycle activity, (ii) uncoupling metabolic and proteostasis remodeling from normal developmental timing, and (iii) inverting the temporal regulation of biosynthetic and metabolic programs between P1 and P7. Specifically, pathways that normally decline during postnatal maturation are aberrantly upregulated in mutants, whereas those that should increase remain suppressed. This mis-timed developmental reprogramming is most pronounced during the P1–P7 window, a critical period when cardiomyocytes normally exit the cell cycle and undergo metabolic and structural maturation.

### Early postnatal repression of Prdm16 accompanies activation of a hypertrophic transcriptional program in Mybpc3 p.Pro109Serfs*12 hearts

To determine whether the transcriptional alterations observed in Mybpc3 p.Pro109Serfs*12 hearts were associated with activation of a hypertrophic gene program, we analyzed the expression of canonical hypertrophy-associated genes at E15.5, P1, P7, and adulthood (Figure 4B). At E15.5, mutant hearts exhibited minimal changes, indicating that overt hypertrophic transcriptional remodeling is not established prenatally (Figure 4B). In contrast, postnatal stages showed progressive activation of a pathological gene program. By P7, classical fetal and stress-responsive markers, including *Nppa, Nppb, Myh7, Ankrd1* and *Acta1*, were significantly upregulated in mutant hearts, with further increases in adulthood, consistent with reactivation of the fetal gene program characteristic of HCM. ^57^ ISH analysis confirmed robust upregulation of *Nppa* and *Ankrd1* in P7 homozygous Mybpc3 p.Pro109Serfs*12 ventricles (Figure 4B).

The transcription factor Prdm16 has been shown to repress the expression of hypertrophic genes in the heart. ^58^ Mechanistically, Prdm16 acts as a negative regulator of MYC target genes, and its downregulation triggers a MYC-dependent cardiac hypertrophic response. ^58, 59^ In addition, *PRDM16* mutations or loss have been associated to both HCM ^60^ and LVNC. ^61, 62^ *Prdm16* transcription was significantly reduced in Mybpc3 p.Pro109Serfs*12 hearts during the postnatal period (Table S3; Figure 4B), coinciding temporally with the emergence of the hypertrophic transcriptional signature. This repression was evident at P7 and persisted into adulthood (Table S3; Figure 4B). Consistent with the marked reduction in *Prdm16* mRNA at P7, we examined whether its downstream targets were transcriptionally altered. Using published whole-heart Prdm16 ChIP-seq data from E13.5 ^63^, we defined a target set of 17,587 binding peaks, including 3,819 promoter-associated peaks (<3 kb from the TSS) corresponding to 3,225 genes. GSEA of these targets against DEGs at each time point showed minimal enrichment in homozygous Mybpc3 p.Pro109Serfs*12 hearts at E15.5. By P1, targets were strongly enriched in mutants, consistent with persistence of an immature gene program. At P7, they became modestly depleted (*p*<0.25), indicating the emergence of a pathological transcriptional state. In adults, a non-significant enrichment trend was observed, consistent with hypertrophic cardiomyopathy–associated immature gene expression (Figure 4C, Figure S17A). To further test how Prdm16 programs shift during the P1-to P7- maturation window, we performed GSEA of the Prdm16 target set across the P1 to P7 transition phase. In the WT P7-P1 comparison, there was no enrichment or depletion of Prdm16 target genes (Figure 4D, Figure S17B), indicating that their expression remains stable across normal postnatal maturation. In contrast, Mybpc3 p.Pro109Serfs*12 (P109S) mutants displayed a marked depletion of Prdm16 target genes in the P7–P1 comparison (P109S P7-P1; Figure 4D, Figure S17B), suggesting that sustained Prdm16-dependent transcriptional programs are required for normal developmental maturation. Together, these findings identify a postnatal window of dysregulated Prdm16 activity that delays maturation before progression toward pathology, consistent with its protective role against hypertrophy and heart failure. ^63^

Given the established role of Prdm16 in the maintenance of ventricular identity, ^63, 64^ cardiomyocyte maturation, ^62^ metabolic regulation, ^65^ and hypertrophic gene expression, ^58, 59^ its downregulation suggested a mechanistic link between impaired maturation programs and pathological remodeling in Mybpc3 p.Pro109Serfs*12 hearts. Thus, the reduction in *Prdm16* expression parallels the pathway-level RNA-seq findings described above (Figure 4A), in which early postnatal stages showed disrupted metabolic and proteostasis programs together with persistent cell cycle enrichment. The temporal association between *Prdm16* downregulation, sustained proliferative signaling, and activation of fetal hypertrophic markers supports a model in which loss of *Mybpc3* perturbs the normal postnatal maturation trajectory, thereby facilitating activation of a pathological hypertrophic program. These data identify *Prdm16* downregulation as a prominent and early molecular feature of Mybpc3-driven HCM, linking defective maturation pathways to progressive hypertrophic remodeling.

### Transgenic Prdm16 expression attenuates the hypertrophic phenotype of Mybpc3 p.Pro109Serfs*12 mice

To investigate the dynamics of Prdm16 expression during ventricular chamber maturation, we performed IF analyses from E16.5 to P7 using a commercially available antibody against human PRDM16 (see *Major Resources Table*). Antibody performance was validated at E18.5 using control and *Prdm16^Δ8^* null mutants (see *Supplemental Methods*). As shown in Figure 4E-E’’, control hearts showed clear nuclear Prdm16 staining, which was absent in *Prdm16^Δ8^* mutants (Figure 4F-F’’), supporting antibody specificity.

At E16.5, Prdm16 expression in control hearts was enriched in the CM relative to the TM (Figure 4G,G’). In contrast, homozygous Mybpc3 p.Pro109Serfs*12 mutants showed increased Prdm16 signal within the TM (Figure 4H,H’,K), consistent with enhanced trabecular growth and hypertrabeculation observed at P1 (Figure 2A-C’’,2N,O,R; Figure S11). At E18.5 and P1, overall Prdm16 distribution appeared comparable between genotypes (Figure S18), despite the enrichment of Prdm16 target gene expression at P1 in Mybpc3 p. P109Sfs*12 mutants (Figure 4C). By P7, IF revealed a reduction of Prdm16 signal in the trabecular/inner myocardium (Figure 4I-J’,K), in agreement with RNA-seq data (Figure 4B) and coinciding with the onset of hypertrophic features in Mybpc3-deficient hearts. Together, these findings indicate that Prdm16 is dynamically regulated during ventricular maturation and suggest that its postnatal downregulation may contribute to the transition from hypertrabeculation to hypertrophy in Mybpc3-deficient hearts.

To assess the functional contribution of Prdm16 to the hypertrophic phenotype observed in Mybpc3 p.Pro109Serfs*12 mutants, we generated a conditional transgenic mouse line in which the mouse *Prdm16* cDNA was inserted into the *Rosa26* locus by homologous recombination in mESC (see *Methods*; Figure S19A,B). In this model, *Prdm16* expression is induced following Cre-mediated excision of a floxed STOP cassette (Figure S19A). We evaluated recombination using the embryonic cardiomyocyte driver Tnnt2^Cre 24^. qRT-PCR confirmed that Cre activation produced a moderate (2x) increase in *Prdm16* transcript levels (Figure S19C).

We next examined whether postnatal transgenic expression of Prdm16 from P1 onwards could rescue the hypertrophic phenotype that emerges in Mybpc3-deficient mice by P7. To this end, we generated Mybpc3 p.Pro109Serfs*12;R26^Prdm16/+^;Myh6^MerCreMer/+^ mice, in which cardiomyocyte specific Cre activity was induced by tamoxifen injection at P1. Whole-mount images of P7 hearts showed that Mybpc3 p.Pro109Serfs*12;R26^Prdm16/+^;Myh6^MerCreMer/+^ hearts were similar in size to control, unlike the enlarged hearts of Mybpc3 p.Pro109Serfs*12 homozygous mutants, although the heart weight to body weight ratio remained comparable to mutants, suggesting a dissociation between gross heart size and relative cardiac mass (Figure 5A,B). Consistently, H&E staining revealed a marked reduction in left ventricular and septal thickness in Prdm16-expressing rescued mice compared with Mybpc3 p.Pro109Serfs*12 homozygotes (Figure 5B,C). IF revealed reduced Prdm16 expression in P7 control hearts (Figure S20A-B’,I), which was further decreased in Mybpc3 p.Pro109Serfs*12 hearts (Figure S20E-F’,I). In contrast, Prdm16 expression was increased in Mybpc3+/+;R26^Prdm16/+^;Myh6^MerCreMer/+^ and, notably, in Mybpc3 p.Pro109Serfs*12;R26^Prdm16/+^;Myh6^MerCreMer/+^ hearts (Figure S20C-D’,G-H’,I). qRT-PCR analysis confirmed a modest (∼2-fold) upregulation of *Prdm16* transcripts in P7 transgenic hearts (Figure S20J). ISH for *Nppa* at P7 revealed robust upregulation in both homozygous Mybpc3 p.Pro109Serfs*12 mutant and Mybpc3 p.Pro109Serfs*12;R26^Prdm16/+^;Myh6^MerCreMer/+^ hearts (Figure 5D), indicating activation of a fetal stress gene program in both genotypes. However, morphometric analysis revealed that cardiomyocyte cross-sectional area was significantly increased in homozygous Mybpc3 p.Pro109Serfs*12 mutants, consistent with cellular hypertrophy, whereas it remained comparable to controls in Mybpc3 p.Pro109Serfs*12;R26^Prdm16/+^;Myh6^MerCreMer/+^ mice (Figure 5E,F; Figure S21). These findings indicate that restoration of Prdm16 expression attenuates cardiomyocyte hypertrophic growth in Mybpc3 p.Pro109Serfs*12 mice, despite persistent activation of fetal gene markers, functionally uncoupling stress-associated transcriptional responses from structural hypertrophy.

**Figure 5.**
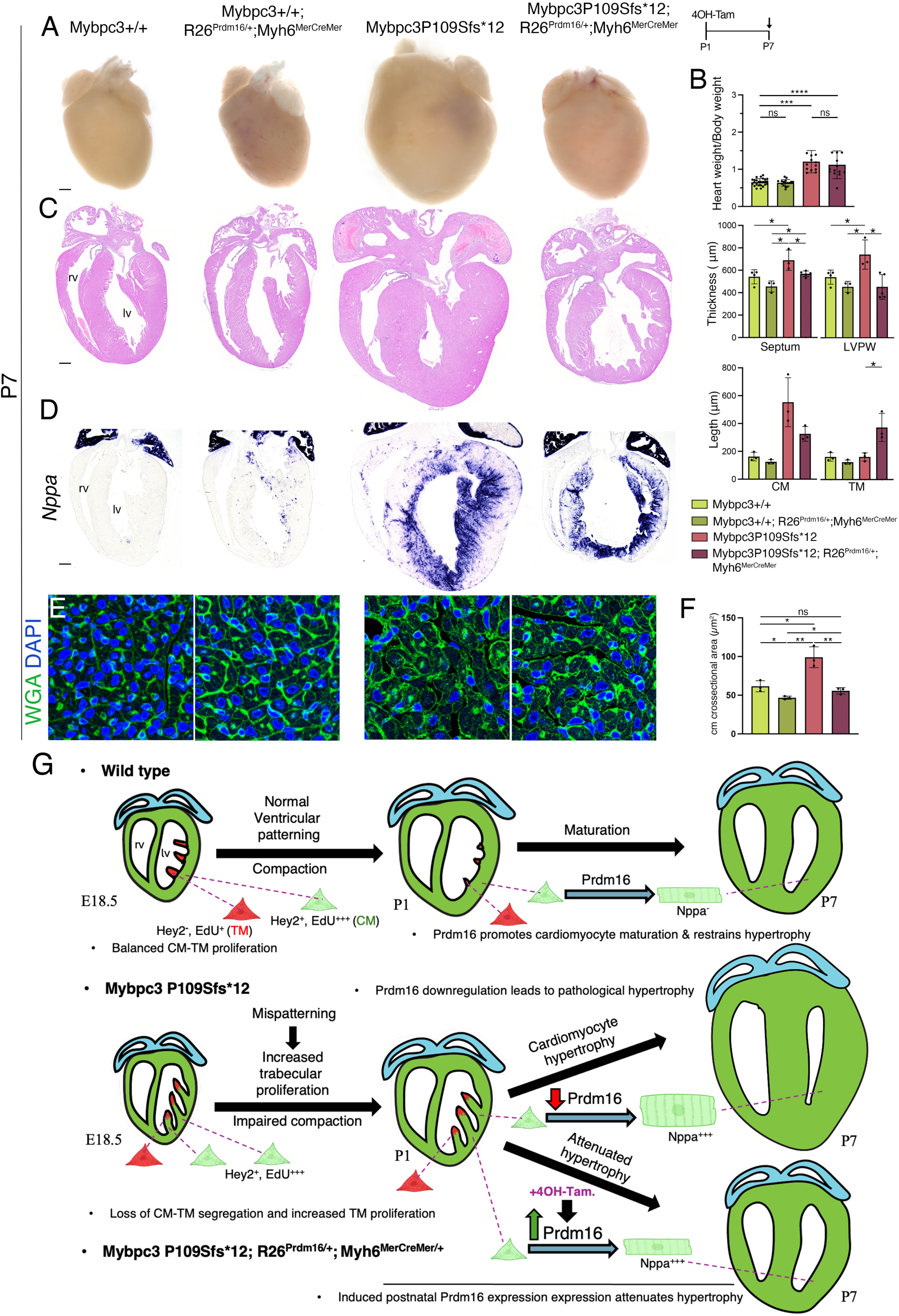
Postnatal cardiomyocyte-specific restoration of Prdm16 attenuates hypertrophy in Mybpc3 p.Pro109Serfs*12 mutant hearts. (**A**) Whole-mount views of P7 control, Mybpc3^+/+^;R26^Prdm16/+^;Myh6^MerCreMer/+^, homozygous Mybpc3 p.Pro109Serfs*12 and Mybpc3 p.Pro109Serfs*12;R26^Prdm16/+^;Myh6^MerCreMer/+^ hearts. Right, Cre-mediated transgenic expression was induced upon Tamoxifen administration at P1. Scale bar, 250 μm. (**B**) Quantification of Heart weight/Body weight ratio (n=25 Mybpc3+/+, n=16 Mybpc3+/+;R26^Prdm16/+^;Myh6^MerCreMer/+^, n=13 homozygous Mybpc3 p.Pro109Serfs*12 and n=14 Mybpc3 p.Pro109Serfs*12;R26^Prdm16/+^;Myh6^MerCreMer/+^), Septum thickness (μm), Left Ventricular Free Wall (LVFW) thickness (μm)(n=4 Mybpc3+/+, n=3 Mybpc3+/+;R26^Prdm16/+^;Myh6^MerCreMer/+^, n=3 homozygous Mybpc3 p.Pro109Serfs*12 and n=5 Mybpc3 p.Pro109Serfs*12;R26^Prdm16/+^;Myh6^MerCreMer^ ≥3 sections per heart), compact myocardium leght (CM) and trabecular myocardium leght (TM) (μm) (n=3 hearts per genotype, ≥3 sections per heart). Data are represented as means ± SD. Statistical significance was determined by unpaired Student’s t-test (ns, not significant; \**p*-value < 0.05; \*\*\**p*-value < 0.001; \*\*\*\**p*-value < 0.0001). (**C**) H&E images of representative hearts of the different genotypes. Scale bar, 250 μm. (**D**) ISH for *Nppa* in the different genotypes. Scale bar, 250 μm. (**E**) WGA (green) and DAPI (blue) fluorescence staining of P7 control, Mybpc3^+/+^;R26^Prdm16/+^;Myh6^MerCreMer/+^, Mybpc3 p.Pro109Serfs*12 and Mybpc3 p.Pro109Serfs*12;R26^Prdm16/+^;Myh6^MerCreMer/+^ heart sections. Scale bar, 10 μm. (**F**) Cardiomyocyte cross sectional area quantification of P7 hearts (n=3, ≥6 sections per heart). Data are represented as means ± SD. Statistical significance was determined by unpaired Student’s t-test (ns, not significant; \**p*-value < 0.05; \*\**p*-value < 0.005). (**G**) Ventricular wall mispatterning drives MYBPC3-associated HCM that is attenuated by Prdm16 expression. Top, wild type: During late gestation (E18.5), compact (CM, Hey2⁺) and trabecular cardiomyocytes (TM, Hey2^-^) are spatially segregated and exhibit balanced proliferation, enabling ventricular compaction progression at birth (P1) and subsequent maturation. Postnatally, Prdm16 promotes cardiomyocyte maturation and restrains hypertrophy, resulting in normal ventricular architecture by P7. Middle, Mybpc3 P109Sfs*12 mutant. Mybpc3 loss disrupts ventricular wall patterning, leading to loss of compact–trabecular segregation, increased trabecular proliferation, and impaired compaction. This is accompanied by peak transcriptional dysregulation from P1 to P7. Abnormally reduced *Prdm16* expression impairs maturation and contributes to pathological cardiomyocyte hypertrophy. Bottom, Mybpc3 P109Sfs*12; R26^Prdm16/+^; Myh6^MerCreMer^ model. Tamoxifen-mediated Prdm16 expression at P1 attenuates hypertrophy despite persistent early patterning defects, promoting cardiomyocyte maturation and partially normalizing ventricular structure.

## DISCUSSION

Genetic cardiomyopathies have classically been diagnosed using imaging and morphological criteria alongside clinical features. Although advances in genomic technologies over the past decade have markedly improved the identification of disease-associated genetic variants, establishing direct genotype–phenotype relationships remains challenging, in part due to substantial phenotypic heterogeneity and overlapping clinical presentations among cardiomyopathy subtypes. ^2, 17^ As a result, functional validation of candidate genetic variants using in vitro or in vivo humanized models remains essential for accurate disease modeling, mechanistic interpretation, and assessment of potential therapeutic strategies. In this context, our findings define a developmental framework through which pathogenic *MYBPC3* variants give rise to distinct cardiomyopathy phenotypes.

Our data show that mice harboring homozygous Mybpc3 p.Pro109Serfs*12 and Mybpc3 p.Arg887Alafs*160 truncating mutations associated with mixed HCM and LVNC in heterozygous human carriers, develop hypertrabeculation/non-compaction during fetal and early postnatal stages. During this developmental window, hypertrabeculation is evident, in the absence of cardiac hypertrophy. Notably, this phenotype is transient and no longer detectable by P7. In contrast, coincident with the resolution of hypertrabeculation, cardiac hypertrophy becomes readily apparent at P7, accompanied by robust upregulation of hypertrophic markers. Together, these observations suggest that hypertrabeculation/LVNC represents a transient developmental state that precedes the onset of HCM following *Mybpc3* abrogation. Consistent with this interpretation, once mutant hearts become hypertrophic, features of hypertrabeculation/LVNC are no longer observed. Moreover, adult mutant mice exhibit sarcomere disarray, altered mitochondrial morphology, and myocardial fibrosis, all hallmarks of HCM. This adult phenotype contrasts with the human condition, in which *MYBPC3* mutation carriers may present with HCM, LVNC, or both. This species difference likely reflects the rapid pace of cardiac maturation in mice, whereby Mybpc3 deficiency manifests as a transient LVNC-like morphology during fetal or early neonatal stages before rapid transition to postnatal hypertrophic remodeling.

Our findings identify the early postnatal period as a critical window during which Mybpc3 loss profoundly disrupts cardiomyocyte transcriptional maturation and metabolic remodeling, thereby predisposing the heart to HCM. While embryonic and adult stages exhibited relatively modest transcriptional differences, the P1-P7 interval showed the most pronounced pathway dysregulation, underscoring the importance of this developmental transition for disease initiation. ^66^ At birth, Mybpc3 p.Pro109Serfs*12 hearts inappropriately maintained strong activation of cell cycle programs, indicating delayed exit from proliferative states that normally accompany cardiomyocyte maturation. This imbalance disproportionately affected the trabecular myocardium, supporting the concept that impaired regulation of cardiomyocyte cell cycle exit contributes to the hypertrabeculated/LVNC-like phenotype. Mechanistically, aberrant activation of TGFβ signaling emerges as a plausible upstream driver of these early defects. TGFβ has been linked to pro-EMT and pro-proliferative programs in the heart ^67^ and implicated in LVNC pathogenesis ^49^, suggesting that inappropriate persistence of this pathway may interfere with ventricular wall maturation during this critical window. Concomitantly, pathways governing oxidative metabolism, mitochondrial function, and translational control were suppressed, revealing an uncoupling of proliferative and metabolic maturation at a stage when cardiomyocytes should rapidly acquire oxidative and biosynthetic competence.

By P7, this imbalance progressed into maladaptive cardiac maturation, with proliferative programs remaining elevated while metabolic and proteostatic pathways became aberrantly upregulated, coincident with overt hypertrophic remodeling. Activation of pathways linked to cell growth, metabolic reprogramming, and oxidative stress is consistent with established molecular hallmarks of HCM. This divergence from the normal maturation trajectory indicates that Mybpc3 loss disrupts the developmental timing of cardiomyocyte growth, metabolism, and RNA surveillance, rather than their absolute activity levels. Over time, this mistimed transcriptional reprogramming culminated in adult hearts with reduced oxidative metabolism and enhanced ECM organization and myogenic programs, consistent with pathological hypertrophic remodeling. Together, these findings support a model in which early defects in cardiomyocyte maturation and proliferative restraint set the stage for maladaptive postnatal remodeling, providing a mechanistic link between developmental hypertrabeculation/LVNC and later HCM. Thus, Mybpc3-related cardiomyopathy arises not solely from adult maladaptive responses, but from an early postnatal failure to correctly execute the transcriptional and metabolic programs that define normal cardiomyocyte maturation.

Although Mybpc3-deficient hearts appeared morphologically normal at midgestation, transcriptomic analyses at E15.5 revealed early alterations in pathways governing protein homeostasis, growth signaling, and metabolism, together with repression of ECM organization and morphogenetic signaling programs. These molecular changes coincide with the onset of ventricular compaction, suggesting that *Mybpc3* loss perturbs the intrinsic maturation state of cardiomyocytes and their surrounding niche at a critical developmental window. Functionally, this dysregulation results in sustained trabecular growth accompanied by aberrant ventricular patterning, marked by ectopic expansion of the Hey2⁺ compact myocardial lineage within the trabecular layer. Lineage-tracing analyses indicate that loss of Mybpc3 disrupts the normal confinement of Hey2⁺-lineage cardiomyocytes to the compact myocardium, leading to their continued incorporation into trabeculae during fetal and early postnatal stages, followed by a relative depletion at later postnatal time points. Importantly, comparable patterning defects were observed in both truncating and, to a lesser extent, missense humanized *Mybpc3* mutations, including the late-onset HCM–associated Mybpc3 p.Arg502Trp allele ^26^, supporting the concept that impaired ventricular patterning constitutes a shared, mutation-independent pathogenic mechanism. In line with this idea, mice carrying humanized LVNC-causing *Mib1* mutations exhibited comparable patterning defects and excessive proliferation,^50, 68^ indicating that altered ventricular patterning and proliferation dynamics are common features of both LVNC and HCM. Together, these findings reposition Mybpc3-related cardiomyopathy as, in part, a disorder of developmental ventricular maturation, in which early perturbations of compact–trabecular lineage dynamics predispose the myocardium to a hypertrabeculation/LVNC-to-HCM transition later in life.

We identify the early postnatal downregulation of *Prdm16* as a key molecular event linking defective cardiomyocyte maturation to pathological hypertrophic remodeling in Mybpc3 - deficient hearts. While prenatal stages showed minimal activation of hypertrophic transcriptional programs, Mybpc3 p.Pro109Serfs*12 hearts underwent a rapid postnatal transition characterized by robust induction of fetal and stress-responsive genes, including *Nppa*, *Nppb*, *Myh7*, *Ankrd1*, and *Acta1*. This transition coincided with a reduction in Prdm16 expression beginning in the early neonatal period and persisting into adulthood, positioning *Prdm16* downregulation as an early feature of Mybpc3-driven cardiomyopathy (Figure 5G). Given the established roles of Prdm16 in maintaining ventricular identity, ^63, 64^ coordinating metabolic maturation, ^65^ and restraining MYC-dependent growth and hypertrophic gene expression, ^58^ its downregulation provides a mechanistic connection between the altered postnatal transcriptional landscape and the emergence of structural hypertrophy. Notably, Prdm16 showed stage-specific and regionally restricted regulation during ventricular maturation: Ectopic trabecular expression at E16.5 may drive excessive trabecular expansion, whereas postnatal loss in the inner myocardium by P7 coincided with hypertrophic remodeling in Mybpc3-deficient mutants, including c-Myc, driving a pre-hypertrophic transcription program normally repressed by Prdm16. ^58^ Importantly, modest postnatal restoration of Prdm16 expression was sufficient to attenuate cardiomyocyte hypertrophy in *Mybpc3* mutants despite persistent activation of fetal stress markers, functionally dissociating stress-associated transcriptional responses from pathological cellular growth. Together, these findings support a model in which Prdm16 acts as a postnatal antihypertrophic regulator downstream of normal sarcomeric function (Figure 5G). Loss of Mybpc3 disrupts normal cardiomyocyte maturation during the early postnatal period, impairing the transition toward a mature contractile state. This maturation defect is accompanied by down-regulation of *Prdm16*, which normally restrains hypertrophic transcriptional programs during this stage. Reduced Prdm16 expression permits premature activation of hypertrophic gene expression, leading to progressive cardiomyocyte enlargement. Thus, defective sarcomeric maturation represents the initiating event, placing *Prdm16* downstream of Mybpc3-dependent contractile maturation. Consequently, early postnatal failure of sarcomere maturation provides a mechanistic link between primary sarcomeric dysfunction and the later emergence of HCM.

## Acknowledgements

We thank the CNIC Transgenesis, Histology, Advanced Imaging, Microscopy and Genomics Units for technical support. We thank ReDIB ICTS infrastructure TRIMA@CNIC, MICIU for the Leica SP8 Navigator microscope. We thank L. Méndez for mouse husbandry.

## Supplemental Material

Supplemental Methods

Figures S1-S21

Tables S1-S4

Major Resources table

ARRIVE checklist for animal studies

## Non-standard Abbreviations and Acronyms

CHD: Congenital heart disease
CM: Compact myocardium
CMRI: Cardiac magnetic resonance imaging
ECHO: echocardiography
ECM: Extracellular matrix
GSEA: Gene set enrichment analysis
HCM: Hypertrophic cardiomyopathy
IF: immunofluorescence
ISH: in situ hybridization
LVNC: Left ventricular non-compaction
MYBPC3: Myosin binding protein c3
mESC: Mouse embryonic stem cells
NMD: Nonsense mediated decay
P1: Postnatal day 1
P7: Postnatal day 7
PTC: Premature termination codon
qRT-PCR: Real time quantitative reverse transcription PCR
ssODN: Single-stranded donor oligonucleotides
TM: Trabecular myocardium
WB: Western blot

## REFERENCES

1. Watkins H, Ashrafian H and Redwood C. Inherited cardiomyopathies. The New England journal of medicine. 2011;364:1643–56.

2. Parikh VN, Day SM, Lakdawala NK, Adler ED, Olivotto I, Seidman CE and Ho CY. Advances in the study and treatment of genetic cardiomyopathies. Cell. 2025;188:901–918.

3. Mazzarotto F, Hawley MH, Beltrami M, Beekman L, de Marvao A, McGurk KA, Statton B, Boschi B, Girolami F, Roberts AM, Lodder EM, Allouba M, Romeih S, Aguib Y, Baksi AJ, Pantazis A, Prasad SK, Cerbai E, Yacoub MH, O’Regan DP, Cook SA, Ware JS, Funke B, Olivotto I, Bezzina CR, Barton PJR and Walsh R. Systematic large-scale assessment of the genetic architecture of left ventricular noncompaction reveals diverse etiologies. Genet Med. 2021;23:856–864.

4. Towbin JA, Lorts A and Jefferies JL. Left ventricular non-compaction cardiomyopathy. Lancet. 2015;386:813–25.

5. Marian AJ and Braunwald E. Hypertrophic Cardiomyopathy: Genetics, Pathogenesis, Clinical Manifestations, Diagnosis, and Therapy. Circ Res. 2017;121:749–770.

6. Jenni R, Oechslin EN and van der Loo B. Isolated ventricular non-compaction of the myocardium in adults. Heart. 2007;93:11–5.

7. Casanova JD, Carrillo JG, Jimenez JM, Munoz JC, Esparza CM, Alvarez MS, Escriba R, Milla EB, de la Pompa JL, Raya A, Gimeno JR, Molina MS and Garcia GB. Trabeculated Myocardium in Hypertrophic Cardiomyopathy: Clinical Consequences. J Clin Med. 2020;9.

8. Hoedemaekers YM, Caliskan K, Michels M, Frohn-Mulder I, van der Smagt JJ, Phefferkorn JE, Wessels MW, ten Cate FJ, Sijbrands EJ, Dooijes D and Majoor-Krakauer DF. The importance of genetic counseling, DNA diagnostics, and cardiologic family screening in left ventricular noncompaction cardiomyopathy. Circ Cardiovasc Genet. 2010;3:232–9.

9. van Waning JI, Caliskan K, Hoedemaekers YM, van Spaendonck-Zwarts KY, Baas AF, Boekholdt SM, van Melle JP, Teske AJ, Asselbergs FW, Backx A, du Marchie Sarvaas GJ, Dalinghaus M, Breur J, Linschoten MPM, Verlooij LA, Kardys I, Dooijes D, Lekanne Deprez RH, AS IJ, van den Berg MP, Hofstra RMW, van Slegtenhorst MA, Jongbloed JDH and Majoor-Krakauer D. Genetics, Clinical Features, and Long-Term Outcome of Noncompaction Cardiomyopathy. J Am Coll Cardiol. 2018;71:711–722.

10. Miller RJH, Heidary S, Pavlovic A, Schlachter A, Dash R, Fleischmann D, Ashley EA, Wheeler MT and Yang PC. Defining genotype-phenotype relationships in patients with hypertrophic cardiomyopathy using cardiovascular magnetic resonance imaging. PLoS One. 2019;14:e0217612.

11. Lin Y, Huang J, Zhu Z, Zhang Z, Xian J, Yang Z, Qin T, Chen L, Huang J, Huang Y, Wu Q, Hu Z, Lin X and Xu G. Overlap phenotypes of the left ventricular noncompaction and hypertrophic cardiomyopathy with complex arrhythmias and heart failure induced by the novel truncated DSC2 mutation. Orphanet J Rare Dis. 2021;16:496.

12. Przytula N, Dziewiecka E, Winiarczyk M, Graczyk K, Stepien A and Rubis P. Hypertrophic cardiomyopathy and left ventricular non-compaction: Distinct diseases or variant phenotypes of a single condition? World J Cardiol. 2024;16:496–501.

13. Harris SP, Bartley CR, Hacker TA, McDonald KS, Douglas PS, Greaser ML, Powers PA and Moss RL. Hypertrophic cardiomyopathy in cardiac myosin binding protein-C knockout mice. Circ Res. 2002;90:594–601.

14. Carrier L, Knoll R, Vignier N, Keller DI, Bausero P, Prudhon B, Isnard R, Ambroisine ML, Fiszman M, Ross J, Jr., Schwartz K and Chien KR. Asymmetric septal hypertrophy in heterozygous cMyBP-C null mice. Cardiovasc Res. 2004;63:293–304.

15. Calaghan SC, Trinick J, Knight PJ and White E. A role for C-protein in the regulation of contraction and intracellular Ca2+ in intact rat ventricular myocytes. J Physiol. 2000;528 Pt 1:151–6.

16. Captur G, Lopes LR, Patel V, Li C, Bassett P, Syrris P, Sado DM, Maestrini V, Mohun TJ, McKenna WJ, Muthurangu V, Elliott PM and Moon JC. Abnormal cardiac formation in hypertrophic cardiomyopathy: fractal analysis of trabeculae and preclinical gene expression. Circ Cardiovasc Genet. 2014;7:241–8.

17. Walsh R, Offerhaus JA, Tadros R and Bezzina CR. Minor hypertrophic cardiomyopathy genes, major insights into the genetics of cardiomyopathies. Nat Rev Cardiol. 2022;19:151–167.

18. Helms AS, Thompson AD, Glazier AA, Hafeez N, Kabani S, Rodriguez J, Yob JM, Woolcock H, Mazzarotto F, Lakdawala NK, Wittekind SG, Pereira AC, Jacoby DL, Colan SD, Ashley EA, Saberi S, Ware JS, Ingles J, Semsarian C, Michels M, Olivotto I, Ho CY and Day SM. Spatial and Functional Distribution of MYBPC3 Pathogenic Variants and Clinical Outcomes in Patients With Hypertrophic Cardiomyopathy. Circ Genom Precis Med. 2020;13:396–405.

19. van Dijk SJ, Dooijes D, dos Remedios C, Michels M, Lamers JM, Winegrad S, Schlossarek S, Carrier L, ten Cate FJ, Stienen GJ and van der Velden J. Cardiac myosin-binding protein C mutations and hypertrophic cardiomyopathy: haploinsufficiency, deranged phosphorylation, and cardiomyocyte dysfunction. Circulation. 2009;119:1473–83.

20. Walsh R, Buchan R, Wilk A, John S, Felkin LE, Thomson KL, Chiaw TH, Loong CCW, Pua CJ, Raphael C, Prasad S, Barton PJ, Funke B, Watkins H, Ware JS and Cook SA. Defining the genetic architecture of hypertrophic cardiomyopathy: re-evaluating the role of non-sarcomeric genes. Eur Heart J. 2017;38:3461–3468.

21. Ho CY, Day SM, Colan SD, Russell MW, Towbin JA, Sherrid MV, Canter CE, Jefferies JL, Murphy AM, Cirino AL, Abraham TP, Taylor M, Mestroni L, Bluemke DA, Jarolim P, Shi L, Sleeper LA, Seidman CE, Orav EJ and Investigators HC. The Burden of Early Phenotypes and the Influence of Wall Thickness in Hypertrophic Cardiomyopathy Mutation Carriers: Findings From the HCMNet Study. JAMA Cardiol. 2017;2:419–428.

22. Muzumdar MD, Tasic B, Miyamichi K, Li L and Luo L. A global double-fluorescent Cre reporter mouse. Genesis. 2007;45:593–605.

23. Tian X, Li Y, He L, Zhang H, Huang X, Liu Q, Pu W, Zhang L, Li Y, Zhao H, Wang Z, Zhu J, Nie Y, Hu S, Sedmera D, Zhong TP, Yu Y, Zhang L, Yan Y, Qiao Z, Wang QD, Wu SM, Pu WT, Anderson RH and Zhou B. Identification of a hybrid myocardial zone in the mammalian heart after birth. Nat Commun. 2017;8:87.

24. Jiao K, Kulessa H, Tompkins K, Zhou Y, Batts L, Baldwin HS and Hogan BL. An essential role of Bmp4 in the atrioventricular septation of the mouse heart. Genes Dev. 2003;17:2362–7.

25. Sohal DS, Nghiem M, Crackower MA, Witt SA, Kimball TR, Tymitz KM, Penninger JM and Molkentin JD. Temporally regulated and tissue-specific gene manipulations in the adult and embryonic heart using a tamoxifen-inducible Cre protein. Circ Res. 2001;89:20–5.

26. Sen-Martin L, Fernández-Tresancos, A., López-Unzu, M. A., Pathak, D., Ferrarini, A., Labrador-Cantarero, V., Sánchez-Ortiz, D., Pricolo, M.R., Vicente, N., Velázquez-Carreras, D., Sánchez-García, L., Nicolás-Ávila, J. A., Sánchez-Díaz, M., Schlossarek, S., Cussó, L., Desco, M., Villalba-Orero, M., Guzmán-Martínez, G., Calvo, E., Barriales-Villa, R., Vázquez, J., Sánchez-Cabo, F., Hidalgo, A., Carrier, L., Spudich, J. A., Ruppel, K.M., Alegre-Cebollada, J. Broad therapeutic benefit of myosin inhibition in hypertrophic cardiomyopathy. bioRxiv. 2026.

27. Sabater-Molina M, Saura D, Garcia-Molina Saez E, Gonzalez-Carrillo J, Polo L, Perez-Sanchez I, Olmo MDC, Oliva-Sandoval MJ, Barriales-Villa R, Carbonell P, Pascual-Figal D and Gimeno JR. A Novel Founder Mutation in MYBPC3: Phenotypic Comparison With the Most Prevalent MYBPC3 Mutation in Spain. Rev Esp Cardiol (Engl Ed). 2017;70:105–114.

28. Perez-Sanchez I, Sabater-Molina M, Rocamora MEN, Glover G, Escudero F, de Mingo Casado P and Gimeno-Blanes JR. Phenotypic Characterization of a Family With An In-frame Deletion in the DMD Gene and Variable Penetrance. Curr Gene Ther. 2018;18:246–251.

29. Ratti J, Rostkova E, Gautel M and Pfuhl M. Structure and interactions of myosin-binding protein C domain C0: cardiac-specific regulation of myosin at its neck? J Biol Chem. 2011;286:12650–8.

30. Dominic KL, Choi J, Holmes JB, Singh M, Majcher MJ and Stelzer JE. The contribution of N-terminal truncated cMyBPC to in vivo cardiac function. J Gen Physiol. 2023;155.

31. Marston S, Copeland O, Jacques A, Livesey K, Tsang V, McKenna WJ, Jalilzadeh S, Carballo S, Redwood C and Watkins H. Evidence from human myectomy samples that MYBPC3 mutations cause hypertrophic cardiomyopathy through haploinsufficiency. Circ Res. 2009;105:219–22.

32. Moolman JA, Reith S, Uhl K, Bailey S, Gautel M, Jeschke B, Fischer C, Ochs J, McKenna WJ, Klues H and Vosberg HP. A newly created splice donor site in exon 25 of the MyBP-C gene is responsible for inherited hypertrophic cardiomyopathy with incomplete disease penetrance. Circulation. 2000;101:1396–402.

33. Rottbauer W, Gautel M, Zehelein J, Labeit S, Franz WM, Fischer C, Vollrath B, Mall G, Dietz R, Kubler W and Katus HA. Novel splice donor site mutation in the cardiac myosin -binding protein-C gene in familial hypertrophic cardiomyopathy. Characterization Of cardiac transcript and protein. J Clin Invest. 1997;100:475–82.

34. Helms AS, Tang VT, O’Leary TS, Friedline S, Wauchope M, Arora A, Wasserman AH, Smith ED, Lee LM, Wen XW, Shavit JA, Liu AP, Previs MJ and Day SM. Effects of MYBPC3 loss-of-function mutations preceding hypertrophic cardiomyopathy. JCI Insight. 2020;5.

35. Chen PP, Patel JR, Powers PA, Fitzsimons DP and Moss RL. Dissociation of structural and functional phenotypes in cardiac myosin-binding protein C conditional knockout mice. Circulation. 2012;126:1194–205.

36. Ho CY, Lopez B, Coelho-Filho OR, Lakdawala NK, Cirino AL, Jarolim P, Kwong R, Gonzalez A, Colan SD, Seidman JG, Diez J and Seidman CE. Myocardial fibrosis as an early manifestation of hypertrophic cardiomyopathy. The New England journal of medicine. 2010;363:552–63.

37. Sandmann C, Klaassen S, Kaski JP, Norrish G and International Paediatric Hypertrophic Cardiomyopathy Consortium i. Contemporary practice and resource availability for genetic testing in paediatric hypertrophic cardiomyopathy. J Med Genet. 2025;62:528–530.

38. Seeger T, Shrestha R, Lam CK, Chen C, McKeithan WL, Lau E, Wnorowski A, McMullen G, Greenhaw M, Lee J, Oikonomopoulos A, Lee S, Yang H, Mercola M, Wheeler M, Ashley EA, Yang F, Karakikes I and Wu JC. A Premature Termination Codon Mutation in MYBPC3 Causes Hypertrophic Cardiomyopathy via Chronic Activation of Nonsense-Mediated Decay. Circulation. 2019;139:799–811.

39. Vignier N, Schlossarek S, Fraysse B, Mearini G, Kramer E, Pointu H, Mougenot N, Guiard J, Reimer R, Hohenberg H, Schwartz K, Vernet M, Eschenhagen T and Carrier L. Nonsense-mediated mRNA decay and ubiquitin-proteasome system regulate cardiac myosin-binding protein C mutant levels in cardiomyopathic mice. Circ Res. 2009;105:239–48.

40. Torres C, Lima-Martinez MM, Rosa FJ, Guerra E, Paoli M, Iacobellis G, Rodney M, Romero-Vecchione E, Luisa Saadtjian M, Zagala M and Rodney H. [Epicardial adipose tissue and its association to plasma adrenomedullin levels in patients with metabolic syndrome]. Endocrinol Nutr. 2011;58:401–8.

41. Sweeney HL and Hammers DW. Muscle Contraction. Cold Spring Harb Perspect Biol. 2018;10.

42. Captur G, Ho CY, Schlossarek S, Kerwin J, Mirabel M, Wilson R, Rosmini S, Obianyo C, Reant P, Bassett P, Cook AC, Lindsay S, McKenna WJ, Mills K, Elliott PM, Mohun TJ, Carrier L and Moon JC. The embryological basis of subclinical hypertrophic cardiomyopathy. Sci Rep. 2016;6:27714.

43. Farrell E, Armstrong AE, Grimes AC, Naya FJ, de Lange WJ and Ralphe JC. Transcriptome Analysis of Cardiac Hypertrophic Growth in MYBPC3-Null Mice Suggests Early Responders in Hypertrophic Remodeling. Front Physiol. 2018;9:1442.

44. Cannon L, Yu ZY, Marciniec T, Waardenberg AJ, Iismaa SE, Nikolova-Krstevski V, Neist E, Ohanian M, Qiu MR, Rainer S, Harvey RP, Feneley MP, Graham RM and Fatkin D. Irreversible triggers for hypertrophic cardiomyopathy are established in the early postnatal period. J Am Coll Cardiol. 2015;65:560–9.

45. Jiang J, Burgon PG, Wakimoto H, Onoue K, Gorham JM, O’Meara CC, Fomovsky G, McConnell BK, Lee RT, Seidman JG and Seidman CE. Cardiac myosin binding protein C regulates postnatal myocyte cytokinesis. Proc Natl Acad Sci U S A. 2015;112:9046–51.

46. Singh SR, Zech ATL, Geertz B, Reischmann-Dusener S, Osinska H, Prondzynski M, Kramer E, Meng Q, Redwood C, van der Velden J, Robbins J, Schlossarek S and Carrier L. Activation of Autophagy Ameliorates Cardiomyopathy in Mybpc3-Targeted Knockin Mice. Circ Heart Fail. 2017;10.

47. Taylor EN, Hoffman MP, Barefield DY, Aninwene GE, 2nd, Abrishamchi AD, Lynch TLt, Govindan S, Osinska H, Robbins J, Sadayappan S and Gilbert RJ. Alterations in Multi-Scale Cardiac Architecture in Association With Phosphorylation of Myosin Binding Protein-C. J Am Heart Assoc. 2016;5:e002836.

48. Yang Q, Sanbe A, Osinska H, Hewett TE, Klevitsky R and Robbins J. In vivo modeling of myosin binding protein C familial hypertrophic cardiomyopathy. Circ Res. 1999;85:841–7.

49. Kodo K, Ong SG, Jahanbani F, Termglinchan V, Hirono K, InanlooRahatloo K, Ebert AD, Shukla P, Abilez OJ, Churko JM, Karakikes I, Jung G, Ichida F, Wu SM, Snyder MP, Bernstein D and Wu JC. iPSC-derived cardiomyocytes reveal abnormal TGF-beta signalling in left ventricular non-compaction cardiomyopathy. Nat Cell Biol. 2016;18:1031–42.

50. Siguero-Alvarez M, Salguero-Jimenez A, Grego-Bessa J, de la Barrera J, MacGrogan D, Prados B, Sanchez-Saez F, Pineiro-Sabaris R, Felipe-Medina N, Torroja C, Gomez MJ, Sabater-Molina M, Escriba R, Richaud-Patin I, Iglesias-Garcia O, Sbroggio M, Callejas S, O’Regan DP, McGurk KA, Dopazo A, Giovinazzo G, Ibanez B, Monserrat L, Perez-Pomares JM, Sanchez-Cabo F, Pendas AM, Raya A, Gimeno-Blanes JR and de la Pompa JL. A Human Hereditary Cardiomyopathy Shares a Genetic Substrate With Bicuspid Aortic Valve. Circulation. 2023;147:47–65.

51. De Lange WJ, Farrell ET, Hernandez JJ, Stempien A, Kreitzer CR, Jacobs DR, Petty DL, Moss RL, Crone WC and Ralphe JC. cMyBP-C ablation in human engineered cardiac tissue causes progressive Ca2+-handling abnormalities. J Gen Physiol. 2023;155.

52. Moens CB, Stanton BR, Parada LF and Rossant J. Defects in heart and lung development in compound heterozygotes for two different targeted mutations at the N-myc locus. Development. 1993;119:485–99.

53. D’Amato G, Luxan G, del Monte-Nieto G, Martinez-Poveda B, Torroja C, Walter W, Bochter MS, Benedito R, Cole S, Martinez F, Hadjantonakis AK, Uemura A, Jimenez-Borreguero LJ and de la Pompa JL. Sequential Notch activation regulates ventricular chamber development. Nat Cell Biol. 2016;18:7–20.

54. Guo Y and Pu WT. Cardiomyocyte Maturation: New Phase in Development. Circ Res. 2020;126:1086–1106.

55. Maroli G and Braun T. The long and winding road of cardiomyocyte maturation. Cardiovasc Res. 2021;117:712–726.

56. Li F, Wang X, Capasso JM and Gerdes AM. Rapid transition of cardiac myocytes from hyperplasia to hypertrophy during postnatal development. J Mol Cell Cardiol. 1996;28:1737–46.

57. Chien KR, Knowlton KU, Zhu H and Chien S. Regulation of cardiac gene expression during myocardial growth and hypertrophy: molecular studies of an adaptive physiologic response. FASEB J. 1991;5:3037–46.

58. Cibi DM, Bi-Lin KW, Shekeran SG, Sandireddy R, Tee N, Singh A, Wu Y, Srinivasan DK, Kovalik JP, Ghosh S, Seale P and Singh MK. Prdm16 Deficiency Leads to Age-Dependent Cardiac Hypertrophy, Adverse Remodeling, Mitochondrial Dysfunction, and Heart Failure. Cell Rep. 2020;33:108288.

59. Kang JO, Ha TW, Jung HU, Lim JE and Oh B. A cardiac-null mutation of Prdm16 causes hypotension in mice with cardiac hypertrophy via increased nitric oxide synthase 1. PLoS One. 2022;17:e0267938.

60. Delplancq G, Tarris G, Vitobello A, Nambot S, Sorlin A, Philippe C, Carmignac V, Duffourd Y, Denis C, Eicher JC, Chevarin M, Millat G, Khallouk B, Rousseau T, Falcon-Eicher S, Vasiljevic A, Harizay FT, Thauvin-Robinet C, Faivre L and Kuentz P. Cardiomyopathy due to PRDM16 mutation: First description of a fetal presentation, with possible modifier genes. Am J Med Genet C Semin Med Genet. 2020;184:129–135.

61. Arndt AK, Schafer S, Drenckhahn JD, Sabeh MK, Plovie ER, Caliebe A, Klopocki E, Musso G, Werdich AA, Kalwa H, Heinig M, Padera RF, Wassilew K, Bluhm J, Harnack C, Martitz J, Barton PJ, Greutmann M, Berger F, Hubner N, Siebert R, Kramer HH, Cook SA, MacRae CA and Klaassen S. Fine mapping of the 1p36 deletion syndrome identifies mutation of PRDM16 as a cause of cardiomyopathy. Am J Hum Genet. 2013;93:67–77.

62. Sun B, Rouzbehani OMT, Kramer RJ, Ghosh R, Perelli RM, Atkins S, Fatahian AN, Davis K, Szulik MW, Goodman MA, Hathaway MA, Chi E, Word TA, Tunuguntla H, Denfield SW, Wehrens XHT, Whitehead KJ, Abdelnasser HY, Warren JS, Wu M, Franklin S, Boudina S and Landstrom AP. Nonsense Variant PRDM16-Q187X Causes Impaired Myocardial Development and TGF-beta Signaling Resulting in Noncompaction Cardiomyopathy in Humans and Mice. Circ Heart Fail. 2023;16:e010351.

63. Wu T, Liang Z, Zhang Z, Liu C, Zhang L, Gu Y, Peterson KL, Evans SM, Fu XD and Chen J. PRDM16 Is a Compact Myocardium-Enriched Transcription Factor Required to Maintain Compact Myocardial Cardiomyocyte Identity in Left Ventricle. Circulation. 2022;145:586–602.

64. Van Wauwe J, Mahy A, Craps S, Ekhteraei-Tousi S, Vrancaert P, Kemps H, Dheedene W, Donate Puertas R, Trenson S, Roderick HL, Beerens M and Luttun A. PRDM16 determines specification of ventricular cardiomyocytes by suppressing alternative cell fates. Life Sci Alliance. 2024;7.

65. Wang W, Ishibashi J, Trefely S, Shao M, Cowan AJ, Sakers A, Lim HW, O’Connor S, Doan MT, Cohen P, Baur JA, King MT, Veech RL, Won KJ, Rabinowitz JD, Snyder NW, Gupta RK and Seale P. A PRDM16-Driven Metabolic Signal from Adipocytes Regulates Precursor Cell Fate. Cell Metab. 2019;30:174–189 e5.

66. Nixon BR, Williams AF, Glennon MS, de Feria AE, Sebag SC, Baldwin HS and Becker JR. Alterations in sarcomere function modify the hyperplastic to hypertrophic transition phase of mammalian cardiomyocyte development. JCI Insight. 2017;2:e90656.

67. Sorensen DW and van Berlo JH. The Role of TGF-beta Signaling in Cardiomyocyte Proliferation. Curr Heart Fail Rep. 2020;17:225–233.

68. Luxan G, Casanova JC, Martinez-Poveda B, Prados B, D’Amato G, MacGrogan D, Gonzalez-Rajal A, Dobarro D, Torroja C, Martinez F, Izquierdo-Garcia JL, Fernandez-Friera L, Sabater-Molina M, Kong YY, Pizarro G, Ibanez B, Medrano C, Garcia-Pavia P, Gimeno JR, Monserrat L, Jimenez-Borreguero LJ and de la Pompa JL. Mutations in the NOTCH pathway regulator MIB1 cause left ventricular noncompaction cardiomyopathy. Nat Med. 2013;19:193–201.

